# Clonal spheroids capture functional and genetic heterogeneity of head and neck cancer

**DOI:** 10.1101/2024.05.24.595655

**Authors:** Jyoti Pandey, Md. Zubbair Malik, Ritis K Shyanti, Palak Parashar, Praveen K Kujur, Deepali Mishra, Dhanir Tailor, Jee Min Lee, Tejinder Kataria, Deepak Gupta, Hitesh Verma, Sanjay V Malhotra, Suneel Kateriya, Vibha Tandon, Rupesh Chaturvedi, Rana P Singh

**Author notes:** Address for correspondence: Prof. Rana P. Singh,; Prof. Rupesh Chaturvedi,; Prof. Vibha Tandon.

## Abstract

Head and neck cancer squamous cell carcinoma (HNSCC) cells exhibit both structural and functional diversity, making them valuable models for understanding tumor heterogeneity at clinical levels. In this study, we generated single-cell-derived spheroids (SCDS) from HNSCC cell lines and patient tumor cells using scaffold- and non-scaffold-based methods to assess this variability. A distinct structural variability among these SCDS, categorized as hypo- and hyperproliferative spheroids based on size, was observed. Hyperproliferative spheroids demonstrated heightened proliferative and tumorigenic potential and increased sensitivity to cisplatin and radiation, while hypoproliferative spheroids exhibited enhanced migratory capabilities. Single-cell RNA sequencing (scRNA-seq) of hypo- and hyperproliferative spheroids provided insights into the transcriptional landscape of HNSCC cells, validating the observed structural and functional heterogeneities within primary tumors. These functionally and genetically characterized spheroids offer valuable tools for the development of next-generation therapeutics.

**Statement of Significance:** Establishment and characterization of single-cell-derived spheroids from head and neck cancer cells, employing scaffold and non-scaffold materials, demonstrate functional and genetic heterogeneity. Single-cell analysis reveals correlations between genetic diversity and spheroid functionality. These characterized spheroids offer potential for advancing therapeutics development.

## Introduction

HNSCC are epithelial tumor that comprises a diverse group of squamous epithelial cancers arising from the oral cavity, sinonasal tract, nasopharynx, larynx, hypopharynx, and oropharynx (1). It is the seventh most prevalent cancer in the world in terms of incidence and accounts for more than 90% of HNC (2). HNC accounts for 30% of all cancer cases in India. Regardless of clinical stage, HNSCCs are a wide range of highly complex diseases with significant levels of inter- and intra-tumoral heterogeneity (3,4). The mechanics and functional ramifications of HNSCC heterogeneity patterns have proven challenging to understand, and therefore, necessitate substantial follow-up research in different model systems.

HNSCC cell line models are frequently employed to explore regulatory systems and gene expression, thus connecting genomic alterations to clinical responses in patients, crucially aiding in cancer therapeutics. Transcriptomic heterogeneity at the single-cell level within HNSCC cell lines can be a useful surrogate to investigate intertumoral heterogeneity in primary and metastatic tumors. Herein, we employed single-cell derived clonal spheroids using selected HNSCC cell lines in an attempt to identify the wide spectrum of genetic and phenotypic variations in tumors reminiscent of a primary or metastatic HNSCC. Single-cell spheroids can recapitulate the intratumor heterogeneity, which is influenced by interactions within the three-dimensional (3D) spheroid microenvironment and by its genetically diverse yet related sub-clonal population. Cancer stem cells (CSCs) or cells with stem cell-like characteristics within these SCDS can leverage cellular plasticity, enhancing structural and functional heterogeneity (5,6). Studies employing single-cell sequencing have identified common recurrent heterogeneous programs of gene expression, which were found in multiple cell lines and patient tumor tissue and were related to the cell cycle, epithelial-mesenchymal transition, which overlapped substantially with patterns observed in clinical settings (7).

In the current study, we established single-cell spheroids from the three HNSCC cell lines, UMSCC-22B, A-253, FaDu and patient tumor tissue to evaluate structural and functional heterogeneity of the HNC cell population. We then performed integrated transcriptome analysis using scRNA seq to gain insight into gene expression modules in HNSCCs to evaluate structural and functional heterogeneity among the UMSCC-22B cell line populations. Further, the established SCDS model could be used for high-throughput drug screening and to understand the disease progression.

## Methods

### Human HNSCC Cell Lines, Cell Culture, and Reagents

UMSCC-22B cells were purchased from Sigma Aldrich, whereas A-253 and FaDu cells were purchased from ATCC. UMSCC-22B, FaDu, and A-253 cells were cultured in Dulbecco’s Modified Eagle Medium (DMEM, Thermo Fisher Scientific), Eagle’s Minimum Essential Medium (EMEM, Sigma Aldrich), and McCoy’s Medium, respectively, supplemented with 1% penicillin-streptomycin (100 U/mL) and 10% FBS. All cell lines were maintained at 37°C with a 5% CO_2_ humidified incubator and were confirmed mycoplasma-negative.

### Generation of Single-cell Derived Spheroids (SCDS)

The generation of SCDS was carried out using two techniques: scaffold-based method using matrigel (BD Biosciences) and scaffold-free method using Ultra Low Attachment (ULA) plates (Corning). For the ULA plate-based method, single-cell suspension of UMSCC-22B, FaDu, and A-253 cells were serially diluted and seeded at three different dilutions (2, 4 and 128 cells/well) with 50 µL serum-free DMEM/F12 medium (Gibco) supplemented with 10 ng/ml human EGF (R & D systems), 10 ng/ml human FGF (R & D systems), human R-spondin 1 (R & D systems), 10 ng/ml murine Wnt-3a (Peprotech), 10 ng/ml murine Noggin (Peprotech), 1 mM N-Acetyl-L-cysteine (NAC) (Sigma-Aldrich), 2% B27 supplement (Gibco), 1% N_2_ supplement (Gibco), 1% Glutamax (Gibco) and 1% penicillin and streptomycin (Sigma Aldrich) in 96-well ULA plate. UMSCC-22B cells were cultured in both round and flat bottom 96-well ULA plates, whereas FaDu and A-253 cells were cultured in only round bottom 96-well ULA plates to grow SCDS. For matrigel-based method, a similar method was used where single-cell suspension of UMSCC-22B cells at three different dilutions (2, 4, and 128 cells/well) with 50 µL matrigel were seeded into wells of matrigel-coated 96-well plate. Cells were then incubated at 37 °C and 5% CO_2_. Every 2-3 days, 25 μL of fresh culture media was replaced in each well.

### Patient Tissue Processing and Spheroid Culture

Patients head and neck tumor tissue were collected for research purposes following the protocol approved by the Institutional Review Board (IRB) at Jawaharlal Nehru University (JNU) (2017-123), All India Institute of Medical Sciences (AIIMS) (2017-123), and Medanta, The Medicity (MICR-1013/2019). Prior written informed consent was obtained from all patients. Harvested fresh tissue specimens at AIIMS were immediately placed in a transport media on ice and transported to the laboratory at JNU to establish SCDS.

The five patient tumor tissues were used to culture spheroids. The stages, gender, and ages of patients are as follows: T2N0M0/M/47, T3N0M0/M/40, T3N2bM0/F/59, T4aN2bM0/M/41, and T4bN2bM0/M/24. Just afterarrival, tumor tissues were washed twice with cold HBSS (Gibco) containing Primocin 100 μg/mL (InvivoGen), cut into small pieces, and digested with freshly prepared digestion media [200 U/ml collagenase IV (Sigma), 5 mM CaCl₂ and 50 U/ml DNase (Sigma) in HBSS)] for 20 minutes at 37°C with agitation. After washing with fresh HBSS and centrifugation (220 rcf, 4 min), dissociated cells were treated with prewarmed TEG [0.025% trypsin (Himedia), 40 μg/ml EGTA (Sigma) and 10 μg/ml polyvinyl alcohol (Sigma)] for 2 minutes at 37°C. Next, trypsin activity was quenched with cold DMEM media with 1% BSA (Sigma), 10 mM HEPES (Sigma), and 1% penicillin-streptomycin (Sigma). Cells were washed and resuspended in DMEM and filtered using a 40 µm cell strainer to obtain a single-cell suspension. As described above, cells were seeded for spheroid formation in ULA 96-well flat bottom plate.

### Quantification of Spheroid Size

Representative images of SCDS were acquired using a Nikon Eclipse TE2000-S microscope. All images were taken at 10X magnification. Images of SCDS were analyzed to measure the diameter using Image J software (NIH). The volume of the SCDS was measured by using the by formula 4/3πr^3^ where r stands for the mean radius of the spheroids.

### Migration Assay

Migration index and migratory cells of both cell lines, UMSCC-22B and A-253, were measured on day 11 for hypo- and hyperproliferative spheroids. We used the following formula for calculating the migration index and the number of migratory cells.

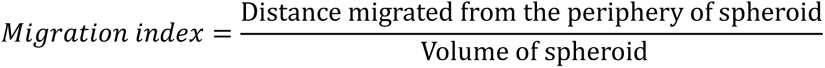

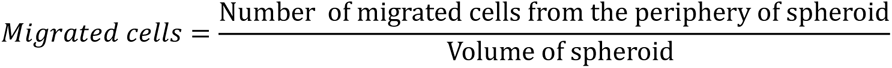

### Confocal Imaging of Spheroids

Spheroids were washed with 1X Phosphate Buffer Saline (PBS) and stained with cocktail dye Hoechst- 33342 (Invitrogen), Calcein-AM (Invitrogen), and Ethidium bromide (EtBr) (Sigma Aldrich). Twenty five µM Calcein-AM (live, green) and 50 µM EtBr (dead, red) reagents were added to each well and incubated for 30 minutes in a CO_2_ incubator and counterstained with 20 µg/ml Hoechst-33342 (Nucleus, blue). Confocal z-slices of spheroids were acquired every 0.4 μm. The images were obtained at 10X and 20X magnifications using confocal microscopy (Nikon A1R HD).

### Spheroid Treatment with Cisplatin and Radiation

The primary tumor cells were cultured from a 65-year-old male patient diagnosed with tongue squamous cell carcinoma, and single-cell derived hypo-and hyperproliferative spheroids were generated following the protocol described earlier. The treatment regimen began on day 7 after the seeding and consisted of 4 hours of cisplatin exposure at a 29.89µM dose. After incubation, the spheroids were washed twice with 1X PBS and once with DMEM (10% FBS). Immediately after that, the spheroids were irradiated followed by 1 hour re-incubation at 37°C. Spheroids were given a single dose of radiation of 4 Gy using an X-ray irradiator at 225 kV and 15 mA at a 50 SSD with a dose rate of 1.28 Gy/min (8) . This was followed by incubating the spheroids with a cocktail solution of calcein-AM and EtBr at 37°C for 30 minutes. After washing with PBS, spheroids were imaged by a fluorescence microscope (OLYMPUS IX2-SP JAPAN) to determine the viability of cells.

### Athymic Nude Mice and Xenograft Study

All the experimental procedures were performed as per the requirements and guidelines of the Committee for Control and Supervision of Experiments on Animals (CCSEA), Government of India, and approved by the Institutional Animal Ethics Committee (IAEC) of JNU. Female BALB/c nude mice (6-week-old) were transplanted subcutaneously on the left/right flank with 10 days old hypoproliferative (15 spheroids/mouse) and hyperproliferative (10 spheroids/mouse) spheroids mixed in 200 µl serum-free medium (SPM): matrigel mixture (1:1). Mice with implanted hypoproliferative spheroids (n=12) and hyperproliferative spheroids (n=8) were monitored daily. Tumor volumes were measured twice weekly and calculated using the formula: volume (mm^3^) = 0.5 × L × W^2^. Kaplan–Meier survival curve was constructed using GraphPad Prism 9.

### Single-cell Capture, Library Preparation and scRNA-seq

Single-cell capture from hypo-and hyperproliferative spheroids and cDNA library preparation was performed using the BD Rhapsody Express Single-cell analysis system (BD Biosciences) following the manufacturer’s instructions. Briefly, hypo-and hyperproliferative spheroids were dissociated into single-cell suspension enzymatically using TrypLE (Gibco), followed by trypsin-EDTA solution (Himedia).

Freshly isolated single-cells from each sample (n=8) were resuspended in sample buffer (BD Biosciences), with a unique sample tag using the BD™ Human Single-Cell Multiplexing kit (BD Biosciences), and loaded on a single BD Rhapsody™ cartridge for single-cell capture. UMI-barcoded magnetic mRNA capture beads were loaded on the cartridge, cells were lysed, and capture beads were retrieved. Library preparation was done using microbead-captured single-cell transcriptome, BD Rhapsody cDNA Kit (BD Biosciences) and the BD Rhapsody whole-transcriptome amplification kit (BD Biosciences). All quantifications during the procedure were done using Qubit (Thermo Fisher Scientific) and 2100 Bioanalyzer System (Agilent) and finally pooled in the ratio – Spheroid mRNA Library : Sample Tag Library: 2.91% : 97.09%. Pooled libraries were sequenced (150 bp pair-ended) on the Illumina HiSeq X platform (Eurofins Genomics, Bangalore).

### ScRNA-seq Data Pre-processing

The obtained FastQ sequencing files were demultiplexed and mapped to the reference genome GRCh38 using the BD Rhapsody WTA Analysis Pipeline in the cloud-based Seven Bridges Platform (Seven Bridges Genomics). Unique molecular identifiers (UMI) count matrices derived from the Rhapsody pipeline were further analyzed using the R package Seurat v4.0.0 (8). Cells having gene numbers between 200-2500 and <15% mitochondrial counts were used for further analysis. The resulting expression matrices were logz-normalized using Seurat’s default parameters. Top 2000 highly variable genes (HVGs) from the corrected expression matrix were extracted and scaled for further analysis.

### Principal Component Analysis, Clustering and Cell-type Annotation

Within the Seurat workflow, the next principal component analysis (PCA) was performed, and the resulting matrices were integrated to remove the batch effects. Next, Louvain clustering was applied with the principal components and followed by Uniform Manifold Approximation and Projection (UMAP) dimension reduction to project the clustered cells onto a two-dimensional (2D) space (9). For cell-type annotation, the CellKb platform (https://www.cellkb.com/) was used to identify different cell populations.

### Differential Expression Gene (DEG) Analysis

The DEGs were performed using the “FindMarkers” function in Seurat in conjunction with the MAST algorithm (version 1.15.0) between different clusters (10). A gene was considered to be upregulated, if the average natural logarithm of the fold change (logFC) > 0.2 and the adjusted P < 0.01; conversely, genes were considered downregulated, if the fold change logFC < −0.2 and adjusted P < 0.01.

### Gene Ontology Analysis

The GO annotation (9) and KEGG pathway enrichment analysis (9) of DEGs were performed using the Database for Annotation Visualization and Integrated Discovery (DAVID; https://david.ncifcrf.gov/). Statistical significance was attributed to findings with a p-value less than 0.05 for ontology terms retrieved from DAVID. On July 14, 2022, the DAVID was accessed.

## Data Availability

The data in the manuscript was generated by the authors and will be made available upon relevant request by the authors.

## Statistical Analysis

Statistical analysis was performed using GraphPad Prism version 9. Paired student t-tests and 2-way ANOVA were performed for statistical analyses in this study. P ≤ 0.05 (*), P ≤ 0.01 (**), P ≤ 0.001 (***), and P ≤ 0.0001 (****) were considered statistically significant.

## Results

### Establishment and Characterization of SCDS from HNC Cell Lines

During the cancer progression, a single-cell begins to transform into a malignant tumor cell and then forms distinct sub-clones of the tumor through a series of mutations, leading to intratumoral heterogeneity (ITH) (11). SCDS is an emerging tool for studying tumor heterogeneity and therapeutic responses *in vitro*. We generated SCDS by seeding HNSCC cells (A-253, FaDu and UMSCC-22B) directly into 96-well round bottom ULA plates or by embedding cells (UMSCC-22B cells) in matrigel and then seeding them onto a flat bottom 96-well plate. SCDS was generated by seeding cells in both scaffold-free (ULA) and scaffold-based (matrigel) methods at the density of 2, 4, and 128 cells/well using the serial dilution method **(Fig. 1A)**. Upon culture of SCDS by matrigel embedding method using UMSCC-22B cells, we observed two stages in the growth of the spheroids: first, the spheroid initiation phase which was followed by the second phase, the spheroid growth phase. The spheroid initiation phase was considered from day 0 to day 4 after cell seeding. During this phase, single-cells proliferated to grow steadily, initiating the clonal spheroid formation in individual wells. In the spheroid growth phase, which starts from day 5 onward, we observed a more compact spherical structure increasing in size and volume over subsequent days, albeit at different growth rates.

**Figure 1.**
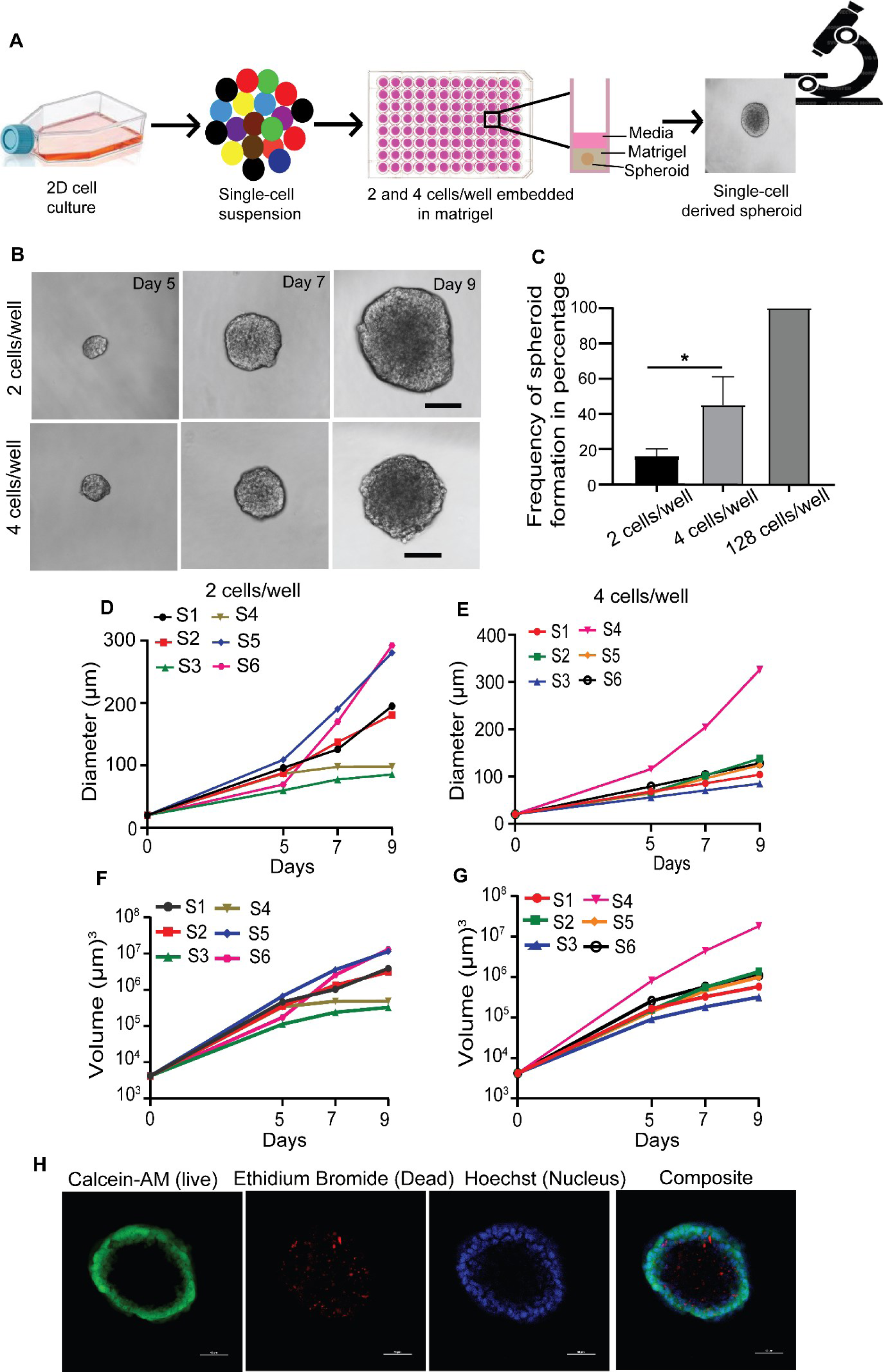
Morphology and growth kinetics of single-cell derived spheroids from UMSCC-22B cells in matrigel. **(A)** Schematic illustration of single-cell derived clonal spheroid generation by matrigel- scaffolding method in ULA microwell plate. **(B)** Representative microscopic images of spheroids from day 5 to day 9 (10 X magnification, scale bar = 100 μm). **(C)** Frequency of spheroids formation from 2, 4, and 128 cells/well. Data are represented as mean ± SEM of three independent experiments (*, P<0.05). **(D-G)** Growth kinetics of spheroids represented by diameter (X and Y planes) and volume (X, Y, and Z planes) (S1-6, Spheroid 1-6). **(H)** Representative confocal images of the single-cell derived spheroid (20 X magnification, scale bar = 50 μm). Calcein-AM (live, green), EtBr (dead, red), and Hoechst-33342 (nuclear, blue) were used for staining.

We next aimed to determine the SCDS formation frequency across different seeding densities in HNC cells. The frequency of SCDS formation was calculated as the percentage of single-cell spheroids successfully formed in each well. The frequency of SCDS formation can be a useful measure of stem cell population within a parent cell population. Generation of SCDS by matrigel embedding method using UMSCC-22B cells at different seeding densities (2, 4, and 128 cells/well) had varied frequencies. The spheroid formation frequency in 128 cells/well was 100 ± 0 % as spheroids formed successfully in each seeded well. A lower frequency of 15 ± 2.6 % was observed in the case of 2 cells/well, whereas 4 cells/well showed a higher frequency, 45 ± 9.2 % **(Fig. 1C).** Similar pattern of spheroid forming frequency was observed when A-253, FaDu, and UMSCC-22B cells were seeded in ULA plates **(Supplementary Fig. S1A-C)**. Additionally, we observed that spheroids were not formed in the wells when seeded with a density of only 1 cell/well. Therefore, we selected cell densities of 2 and 4 cells/well for further work in this study.

Spheroids having a diameter greater than 200 μm were characterized by a heterogeneous population of cells, displaying concentric zones of peripheral proliferating cells, middle quiescent cells, and an inner necrotic core which was apparent as a dark central region (12). With the increase in the size of spheroids of UMSCC-22B cells after day 5, we observed within the spherical geometry of these clonally propagated spheroid an optically dark core region surrounded by strong light-absorbing peripheral region under a light microscope **(Fig. 1B).** Similar demarcation in cellular structure was observed in spheroids formed by UMSCC-22B, FaDu, and A-253 cells when cultured *via* matrigel-free method. Morphology of clonal spheroids formed by UMSCC-22B cells and A-253 (ULA method) had compact morphologies with a defined smooth border. Meanwhile, FaDu cells formed loose aggregate-like structures, implying the lack of proper cell-cell adhesion. **(Supplementary Fig. S2A-B, S2G-H, S2M-N, and S4A).**

Analysis of growth kinetics of spheroid formation revealed structural heterogeneity in terms of size (diameter or volume), which was independent of the initial number of cells seeded. In the matrigel method, the average diameter and the volume of the UMSCC-22B spheroids cultured from 2 and 4 cells/well were 188.6 ± 35.66 µm and 135.7 ± 21.53 µm, respectively, and 5.3 × 10^6^ ± 2.2 × 10^6^ µm^3^ and 3.7 × 10^6^ ± 2.8 × 10^6^ µm^3^, respectively, at day 9. By the seeding density of 2 cells/well the smallest spheroid (diameter- 85.46 µm, volume- 3.26 × 10^5^ µm^3^) and largest spheroid (diameter- 292.24 µm, volume- 1.3 × 10^7^ µm^3^) revealed the variability in the cellular population of UMSCC-22B cells **(Fig. 1D and F)**. Similarly, at the seeding density of 4 cells/well, the spheroid diameter ranged from 84.88 μm to 236.18μm, whereas the spheroid volume ranged from 3.2 × 10^5^ µm^3^ to 1.8 × 10^7^ µm^3^ at day 9 **(Fig. 1E and G)**. Again, a similar trend of spheroid growth was observed with UMSCC-22B,FaDu and A-253 cells when the matrigel-free method was used (**Supplementary Fig. S2C-F, Fig. S2I-L, Fig. S2O-R, Fig. S4B-C)**.

### SCDS Recapitulates Characteristic Spheroid Physiology

As spheroids grow, they closely resemble *in vivo* avascular solid tumors and micrometastases containing heterogeneous cellular zones. These zones, known as proliferating zones present in the periphery, intermediate quiescent zone (slow-dividing cells), and an inner necrotic zone, mimic key elements of the tumor microenvironment, such as oxygen, nutrition pH, and cellular density gradients. Confocal analysis of these SCDS after Hoechst, Calcein-AM, and EtBr staining during the spheroids growing phase at day 10 clearly reveals the outer proliferating zone and inner necrotic zone **(Fig. 1H, Supplementary Fig. S3A-F, Fig. S4D and E).** Our SCDS recapitulated the characteristic features of spheroids, which confirms the biological functionality of these spheroids.

### Hypo- and Hyperproliferative Spheroids Represent Structural Heterogeneity in HNC Cells

Heterogeneity is a vital feature of solid tumors, which can be attributed to differential cellular growth kinetics within clonal subpopulations. Single-cell spheroid formation by both the matrigel-free and matrigel-based methods revealed heterogeneous sizes of spheroids by UMSCC-22B cells. Based on their sizes (diameter), we delineate them into spheroids having hypoproliferative (diameter ≤ 200 µm) and hyperproliferative (diameter > 200 µm) characteristics. The size difference between hypo- and hyperproliferative spheroids was observed clearly on day 11, and we measured their growth in terms of diameter and volume on a reverse day till day 5. A t-test (non-parametric) was carried out to identify differences between those spheroids. A significant difference (P ˂ 0.001 and P ˂ 0.0001) was observed between hypo- and hyperproliferative spheroids (**Supplementary Fig. 5A-E)**. Both hypo- and hyperproliferative spheroids showed a compact morphology and a spherical shape with clear and smooth edges **(Fig. 2A and B, Supplementary Fig. S6A and B)**. Using the matrigel-based method (with a cell density of 2 cells/well) at day 11, the diameter of hypoproliferative spheroids was 150 ± 19.5 µm, and their volume was 2.4 x 10^6^ ± 8.5 x 10^5^ µm^3^. However, the diameter of hyperproliferative spheroids was 367 ± 31.8 µm, and the volume was 3 x 10^7^ ± 8.5 x 10^6^ µm^3^ **(Fig. 2C and 2D).** When UMSCC-22B cells were seeded in a ULA flat bottom plate, similar variations in diameter and volume were observed at a cell density of 2 cells/well (**Supplementary Fig. S6C and D).**

**Figure 2.**
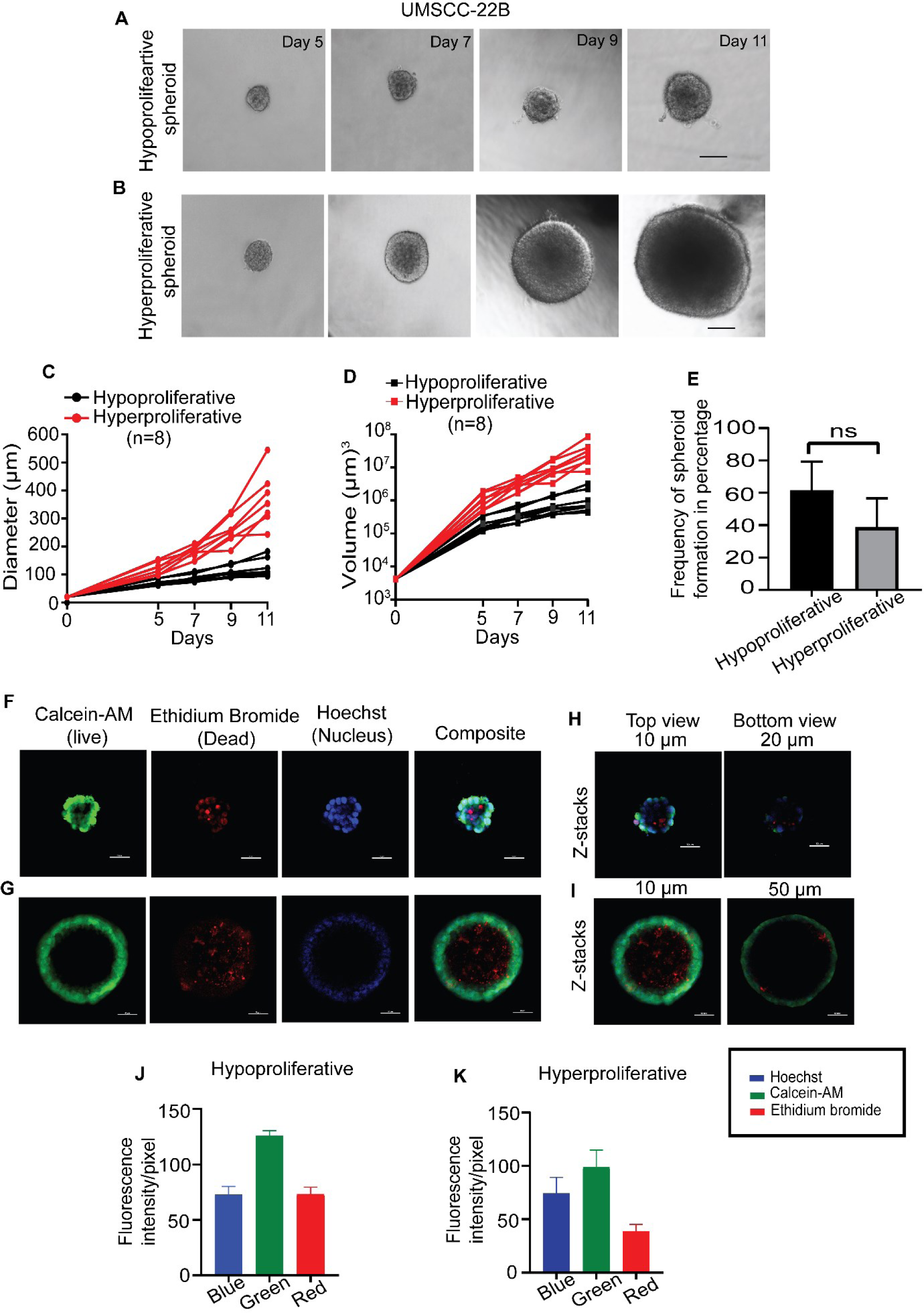
Structural heterogeneity is observed in single-cell derived spheroids UMSCC-22B cells grown in matrigel. **(A and B)** Representative microscopic images of hypo-and hyperproliferative spheroids (10X magnification, scale bar is 100 μm). **(C and D)** Growth kinetics of hypo-and hyperproliferative spheroids in terms of diameter (X and Y planes) and volume (X, Y, and Z planes). **(E)** Frequency of spheroids formation of hypo-and hyperproliferative spheroids. **(F-G)** Representative confocal images of the single-cell derived spheroid. Calcein-AM (live, green), EtBr (dead, red), and Hoechst-33342 (nuclear, blue) were used for staining. **(H-I)** Representative Z-stacked confocal images of hypo-and hyperproliferative spheroids at various Z-planes. **(J-K)** Quantification of fluorescence intensity of dyes and fluorescence intensity/pixel is represented as the mean ± SEM obtained from n=3 spheroids. 20 X magnification, scale bar = 50 μm.

The frequency of hypo- and hyperproliferative spheroids formation is also an important variable in assessing the functional heterogeneity in UMSCC-22B cells. The frequency of hypoproliferative spheroid formation by the matrigel-based method was 61.33 ± 10.35%, whereas the frequency of hyperproliferative spheroid formation decreased to 38.67 ± 10.35% **(Fig. 2E).** Similarly, in the ULA plate (flat bottom), the frequency of hypoproliferative spheroid formation was 53.67 ± 12.45%; whereas, the frequency of hyperproliferative spheroid formation decreased to 46.33 ± 12.45% **(Supplementary Fig. S6E).** Both UMSCC-22B cell line derived hypo- and hyperproliferative spheroids exhibited more Calcein-AM fluorescence intensities (125.3 ± 6.3 and 98.39 ± 9.5, arbitrary unit, respectively) at the periphery region and a comparably lesser EtBr fluorescence intensities (68.4 ± 9 and 38.15 ± 4, respectively) at the core region. Representative focal planes (10 µm and 20 µm for hypo-; 10 µm and 50 µm for hyperproliferative spheroid) from confocal image z-stacks showed similar differences in Calcein-AM/EtBr staining **(Fig. 2F-K, Supplementary Fig. S6F-I, Supplementary video 1)**.

### Patient-derived Spheroids Confirmed Structural Heterogeneity in HNC Cells

Next, we evaluated the clinical concordance of the above growth kinetics studies, using single-cells isolated from the treatment-naive HNC patient tumor (T2N0M0). Single-cell suspension processed directly from the patient tumors was seeded at different densities (25, 50, 100, 200, 500 cells/well) in a ULA flat-bottom plate. Selected spheroids were longitudinally tracked over 9 days to evaluate structural variability among the patient-derived single-cells. As tumor cells isolated from the tumor of HNC patients took a longer time to grow as compared to established cell lines, the patient SCDS was considered hypoproliferative at a diameter ≤ 70 µm and hyperproliferative at a diameter > 70 µm.

Based on the spheroid size and cell proliferation rate, we could clearly segregate the spheroids into hypo- and hyperproliferative spheroids **(Fig. 3A and B, Supplementary Fig. S7A and B)**. Hypoproliferative spheroids had a diameter of 57.34 ± 1.364 µm on day 9, which resulted in a volume of 9.9 x 10^4^ ± 7 x 10^3^ µm^3^. The diameter of hyperproliferative spheroids on day 9 was 113.6 ± 7.6 µm, and their volume was 8 x 10^5^ ± 1.4 x 10^5^ µm^3^ **(Fig. 3C and D, Supplementary Fig. S7C and D).** Diameter and volume analyses depicted the smaller average size of SCDS formed from the HNC patient tumor cells as compared to established HNC cell lines. Irrespective of the seeding density, the frequency of hypoproliferative spheroid formation was always greater than that of hyperproliferative spheroid formation (T2N0M0) **(Fig. 3E)**. We observed the similar staining pattern with higher intensity of Calcein-AM at the rim of the spheroid as compared to the intensities of EtBr at the inner core in HNC patient cells derived hypo- and hyperproliferative spheroids **(Fig. 3F-I)**. These Calcein-AM/EtBr staining patterns implicate towards the structural and functional differences of hypo- and hyperproliferative spheroids, and that may be utilized to unravel intratumoral heterogeneity in HNC patients.

**Figure 3.**
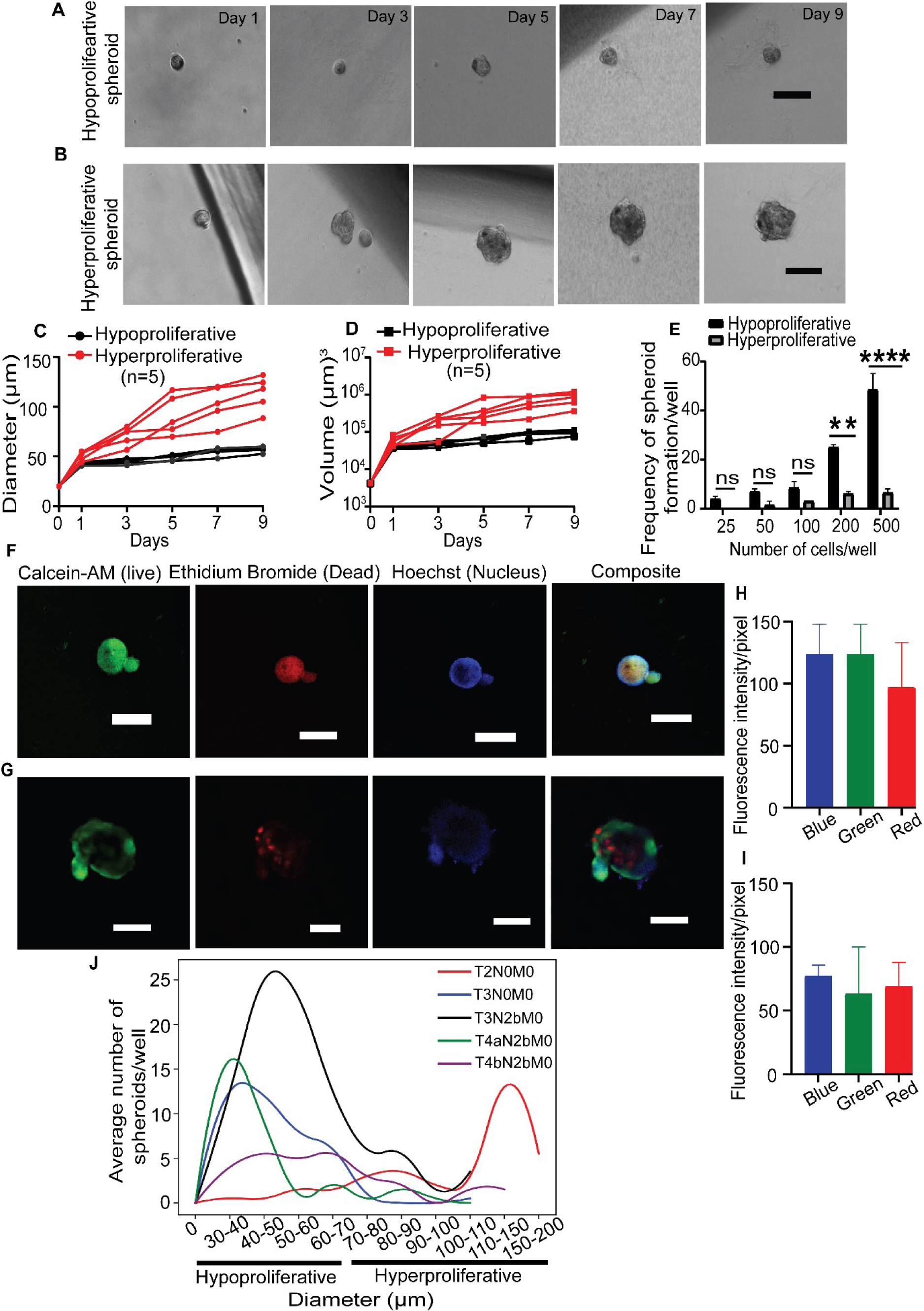
Structural heterogeneity is observed in HNC patient-derived single-cell spheroids in a ULA microplate. **(A and B)** Representative microscopic images of hypo-and hyperproliferative spheroids (10X magnification, scale bar is 100 μm). **(C and D)** Growth kinetics of hypo-and hyperproliferative spheroids in terms of diameter (X and Y planes) and volume (X, Y, and Z planes). **(E)** Frequency of spheroids formation of hypo-and hyperproliferative spheroids. (ns-non-significant, ** p≤0.01, **** p≤0.0001). **(F-G)** Representative confocal images of hypo-and hyperproliferative spheroid. (20X magnification, scale bar = 100 μm). **(H-I)** Quantification of fluorescence intensity of dyes and fluorescence intensity/pixel is represented as the mean ± SEM obtained from n=3 spheroids. **(J)** Frequency of spheroid formation (100 cells/well) from tumors from different stages of HNC patients.

Moreover, we examined the distribution of spheroid sizes in cells isolated from tumors across various TNM stages of head and neck cancer (HNC) patients. Our findings revealed that in early-stage tumors (T2N0M0), hyperproliferative-sized spheroids predominated, while the opposite trend was observed in later-stage tumors (T3N0M0, T3N2bM0, T4aN2bM0, and T4bN2bM0) of HNC patients. **(Fig. 3J)**.

### Hypoproliferative Spheroid Cells Show Higher Migratory Potential than Hyperproliferative Spheroid Cells

During metastatic progression, migratory cells endowed with phenotypic plasticity greatly amplify tumor heterogeneity, affecting therapeutic response in HNC patients. The structural variance between hypo- and hyperproliferative spheroids further encouraged us to evaluate their migration ability. Quantitative measurement of the migratory potential of the spheroids was determined using the migration index, which measured distance travelled by the cells from the outer periphery of the spheroids. In addition, migratory potential was also determined by the count of migratory cells from each spheroid. The migration index of UMSCC-22B-derived hyperproliferative spheroids (1.73 x 10^-5^ ± 3.18 x 10^-6^/µm^2^) was significantly lesser (P ˂ 0.0001) than that of the hypoproliferative spheroids (8.59 x 10^-4^ ± 1.43 x 10^-4^/µm^2^). Similarly, the number of migrated cells in a hyperproliferative spheroid (9.66 x 10^-6^ ± 1.90 x 10^-6^/µm^3^) was lesser than hypoproliferative spheroid (1.21 x 10^-4^ ± 1.89 x 10^-5^/µm^3^) **(Fig. 4A - D)**. We also observed that the migration index and the number of migrated cells in hyperproliferative spheroids were significantly less (P ˂ 0.01) compared to hypoproliferative spheroid in the case of A253 cells (**Supplementary Fig. S8A - D).** These data effectively captured the functional heterogeneity among the SCDS established from UMSCC-22B and A-253 cells.

**Figure 4.**
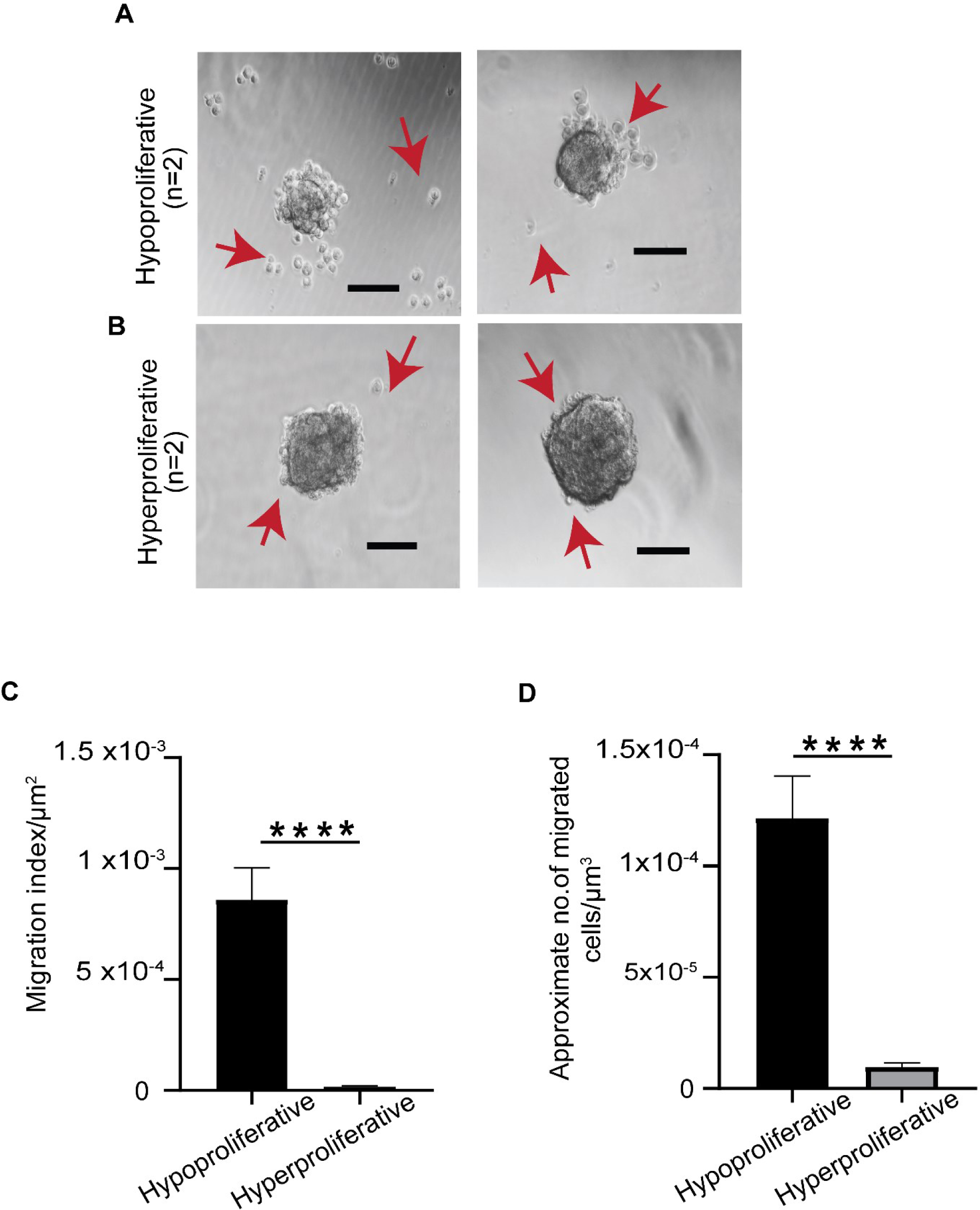
3D single-cell spheroid migration assay reveals functional heterogeneity in hypo-and hyperproliferative spheroids (UMSCC-22B). (**A and B)** Representative images of cell migration from hypo-and hyperproliferative spheroids. (10x magnification, scale bar = 100 μm). **(C)** Quantification of migration using migration index of hypo- and hyperproliferative spheroid. **(D)** Quantification of no. of migrated cells from the periphery of spheroids. (**** p<0.0001).

### Tumor-propagating Potential of Hypo- and Hyperproliferative Spheroids in the Xenograft Model

Further, we evaluated the tumorigenic potential of single-cell derived hypo-and hyperproliferative spheroids *in vivo*, using an athymic nude mice xenograft model. To maintain equivalency in inoculation cell number, we subcutaneously transplanted 15 hypo- and 10 hyperproliferative intact spheroids into mice and evaluated tumor growth over time **(Fig. 5A and B)**. Hypoproliferative spheroids did not form tumor in athymic nude mice (0/12). Contrarily, hyperproliferative spheroids were associated with a 50% (4/8) of tumour incidence **(Fig. 5C).** The tumor appearance in the case of hyperproliferative spheroids (n=4) was noted on days 32, 40, 44, and 52, respectively. **(Fig. 5D)**. Percentage survival for both groups of mice was calculated using the Kaplan-Meier method **(Fig. 5E)**. The median survival of mice xenografted with hypo- and hyperproliferative was 67 and 63 days, respectively, which was found non-significant (Log-rank (Mantel-Cox) test P = 0.5803, χ2=0.3058, and df = 1). These results further suggested that xenograft transplantation did not significantly affect the survival of mice in both groups. However, hyperproliferative spheroids were found to initiate the tumors 50% of mice, whereas hypoproliferative spheroids had no tumorigenic potential.

**Figure 5.**
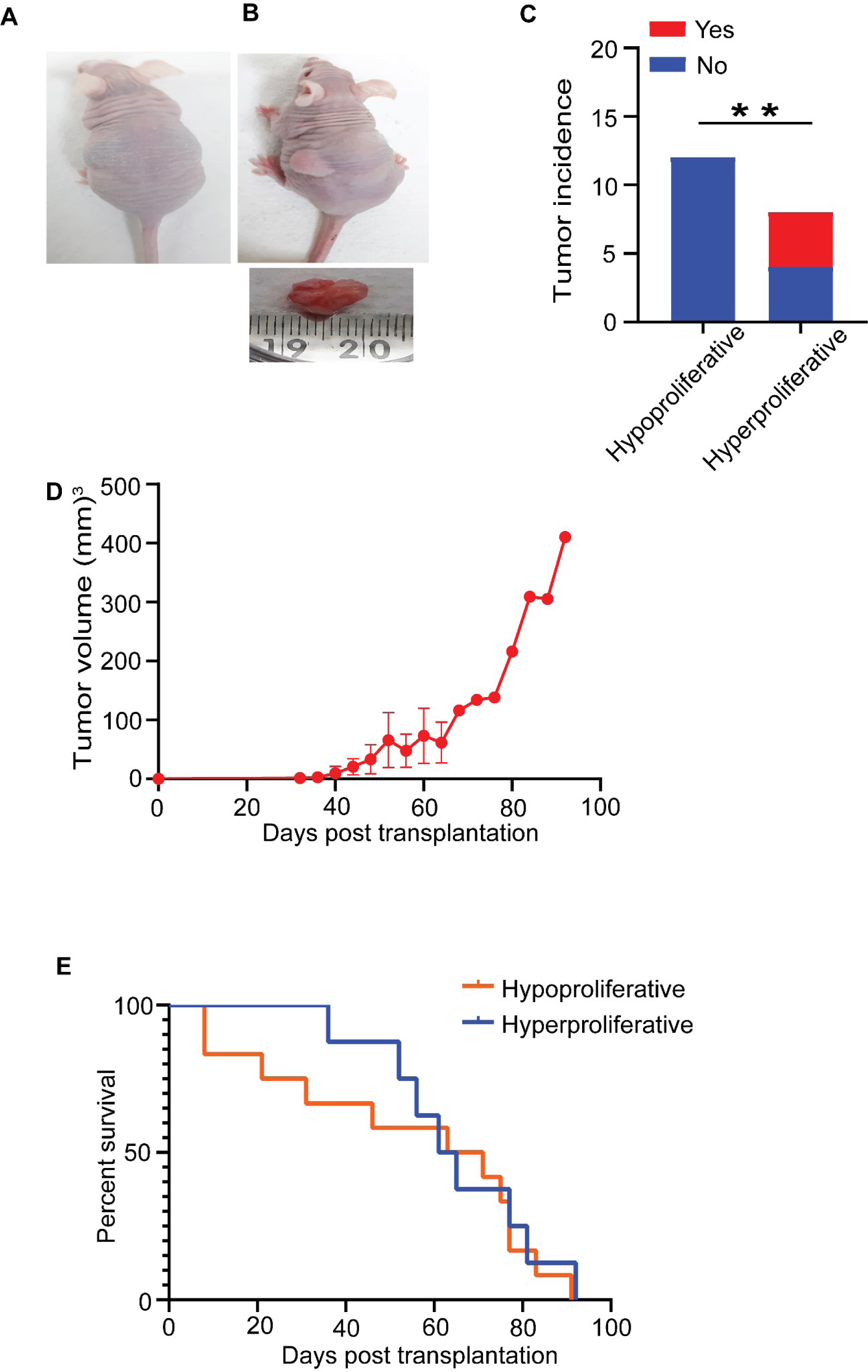
Hyperproliferative spheroids exhibited tumorigenic potential in nude mice. **(A and B)** Representative images of tumorigenesis in athymic nude mice and their corresponding tumor subcutaneously inoculated with hypo-and hyperproliferative spheroids. **(C)** Tumour incidence in athymic nude mice inoculated with hypo-and hyperproliferative spheroids (** p≤0.01). **(D)** Tumour volume after transplantation of hyperproliferative spheroids into athymic nude mice . **(E)** Kaplan-Meier plot of overall survival of athymic nude mice after transplantation with hypo-and hyperproliferative spheroids.

### Effect of Cisplatin and Radiation on Hypo- and Hyperproliferative Spheroids

We further evaluated the efficacy of cisplatin and radiation on hypo- and hyperproliferative spheroids. Hypo- and hyperproliferative spheroids were cultured for 7 days, and on 7^th^ day, these spheroids were exposed to cisplatin, radiation, and a combination of both. Control hypo-and hyperproliferative spheroids appeared to be compact. Individual radiation and cisplatin treatment decreased the compactness of both hypo- and hyperproliferative spheroids, but the combination of both further decreased the compactness of spheroids prominently. The cocktail staining of Calcein-AM and EtBr clearly showed the disintegration of the spheroids due to the treatments **(Fig. 6A and B)**. Normalized fluorescence intensity quantification of calcein-AM and EtBr further showed that the different treatment groups, 4 Gy, cisplatin, and 4Gy + cisplatin, resulted in a significant decrease (P ˂ 0.01 and P ˂ 0.0001) in calcein-AM intensity of hyperproliferative spheroids as compared to hypoproliferative spheroids. While the corresponding EtBr intensity significantly increased (P ≤ 0.05 and P ≤ 0.0001) in hyperproliferative spheroids as compared to hypoproliferative spheroids **(Fig. 6C and D)**. These results confirmed that the hyperproliferative spheroids were more sensitive to cisplatin as well as radiation, and a combination of both than hypoproliferative spheroids.

**Figure 6.**
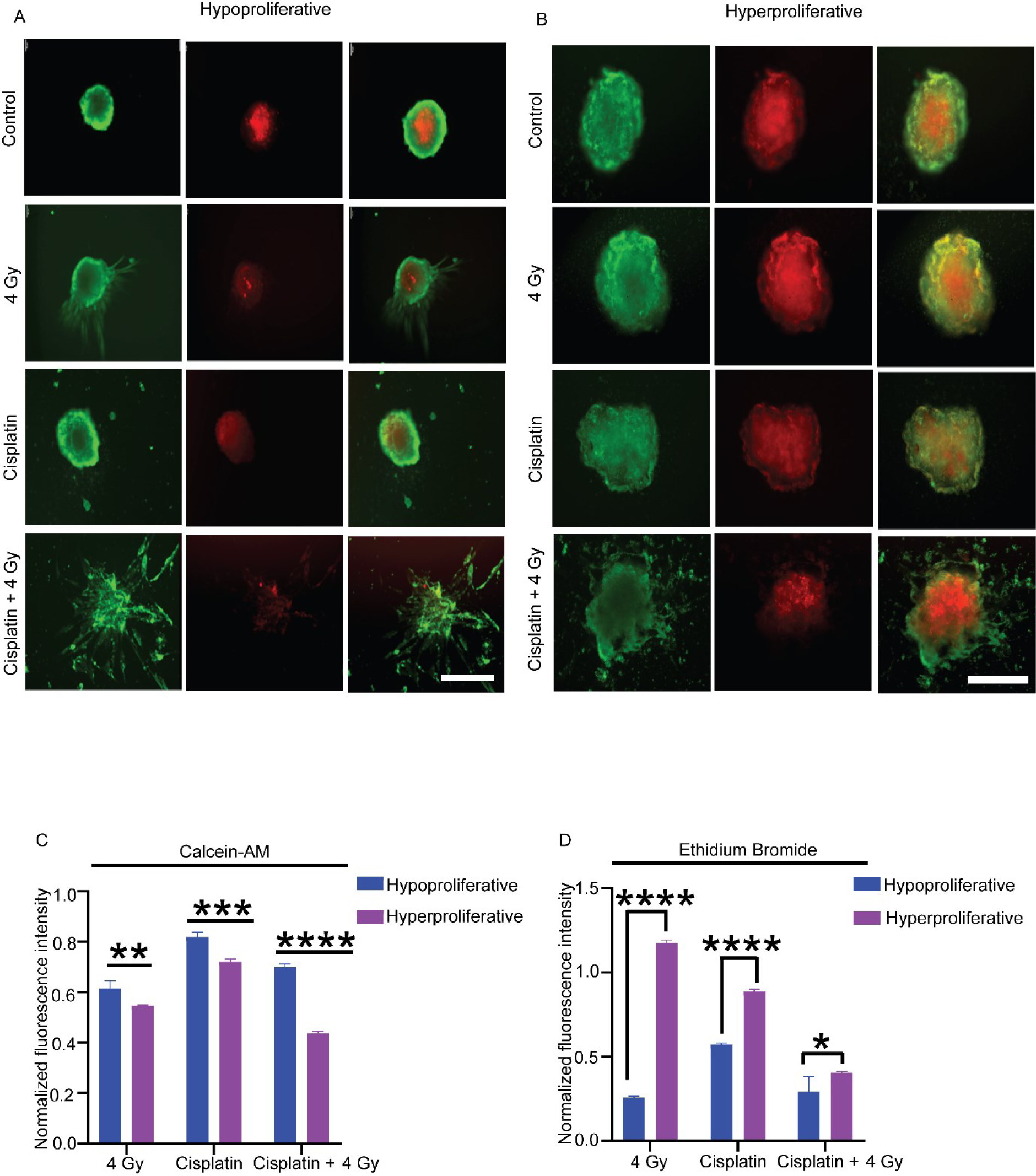
Cell viability of HNC patient-derived spheroids after treatment with cisplatin and radiation. Viable cells appear as green, while non-viable cells appear as red. **(A and B)** Representative fluorescence images of spheroids after treatment with cisplatin, radiation and radiation + cisplatin (scale bar = 100 μm). **(C and D)** Normalized fluorescence intensity of Calcein-AM and Ethidium Bromide for hypo-and hyperproliferative spheroids. (* p<0.05; ** p<0.01, *** p<0.001, **** p<0.0001).

### scRNAseq Analysis, Key Genes, Enriched Pathways and Cell Types in Functional Heterogeneity

To delineate functional heterogeneity among SCDS, a comparative analysis of transcriptomic landscapes of hypo- and hyperproliferative spheroids at the single-cell resolution was done. After the dissociation of spheroids, a total of 10,164 cells were captured, and after quality filtering, 1039 cells (751 cells from hypoproliferative spheroids and 288 cells from hyperproliferative spheroids) were analysed. We filtered out the single-cell transcriptomes with unique features exceeding 2,500 or less than 200 and mitochondrial gene contents of >15%. For a global representation of the cells, we applied the UMAP reduction plot **(Fig. 7A and B)**. We identified 6 distinct clusters (numbered from 0 to 5) in hypo- and 4 distinct clusters (numbered 0 to 3) in hyperproliferative spheroids. Further, DEG analysis was performed for cluster annotation to derive unique DEGs defining the clusters using parameter |log2 fold change| >1.0 with an adjusted P-value < 0.05. Except for clusters C3 and C5 in hypoproliferative spheroids, unique DEGs were identified for each cluster **(Supplementary Fig. 9A and B, Supplementary Tables 1 and 2)**.

**Figure 7.**
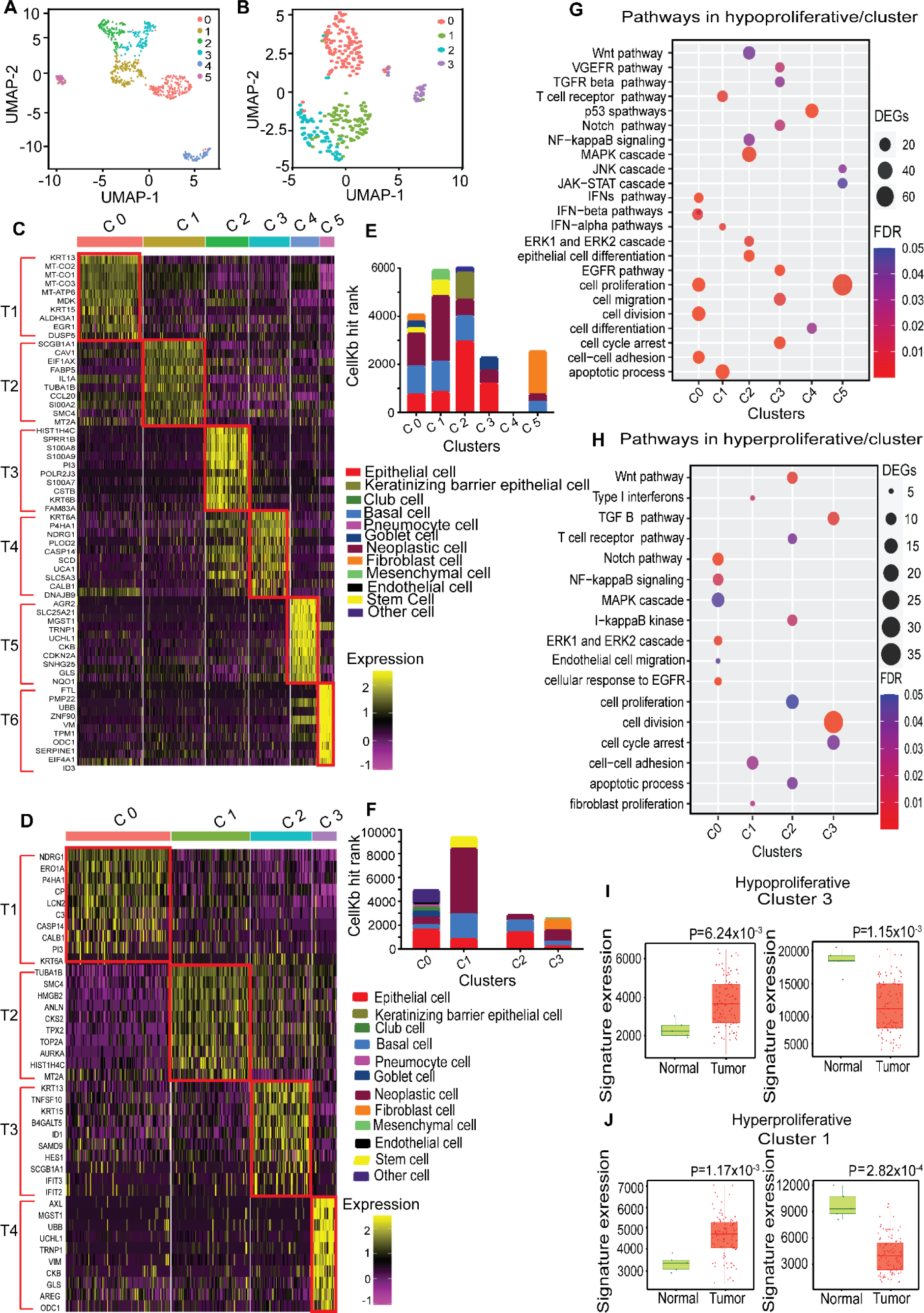
Single-cell transcriptome analysis highlights functional heterogeneity among hypo-and hyperproliferative spheroids. **(A and B)** UMAP visualization of color-coded cell clusters from integrated analysis of single-cell transcriptome data sets from hypo-and hyperproliferative spheroids. **(C and D)** Heatmap of top 10 DEGs for each cluster of hypo-and hyperproliferative spheroids. Cells are represented in columns, and genes are listed in rows. Yellow to dark purple intensities represent high to low gene expression. **(E and F)** Stacked bar plots for cell type annotation in different clusters of hypo-and hyperproliferative spheroids indicate cell-type proportions. The Y-axis represents the CellKb hit rank. **(G and H)** The dot plot for GO analysis showed the foldenrichment of DEGs (size) and -log10 p-value (colour) for selected biological processes per identified clusters in hypo-and hyperproliferative spheroids. **(I and J)** Comparison of scRNA seq data (top ten up and down-regulated genes) with TCGA data. TNM plot was used to determine the expression of top ten up-anddown-regulated genes in normal (3691) and tumor (29,376) HNC. Bar graph depicting the up and downregulated genes in cluster 3 of hypoproliferative and cluster 1 of hyperproliferative spheroid.

The DEG profile was further used to construct the cell-type landscape of these hypo- and hyperproliferative spheroids using the CellKb database. Interestingly, we found different proportions and compositions of cell types among different clusters of hypo-and hyperproliferative spheroids. In the predicted cell types across different clusters, neoplastic, epithelial, and basal cells were in the majority, whereas the minor others were stem, goblet, club, pneumocyte, endothelial, mesenchymal, fibroblast, and keratinizing barrier epithelial and other cells, reflecting the multi-cell type tumor microenvironment in HNC **(Fig. 7E and F, Supplementary Table 5)**.

Heatmap of scaled gene expression of the top ten most expressed genes from each cluster of hypo- and hyperproliferative spheroids identified the expression patterns in coherence with the functional characterization observed in our studies **(Fig. 7C and D, Supplementary Tables 3 and 4)**. We previously observed that hypo- and hyperproliferative spheroids from UMSCC-22B cells have distinct morphological phenotypes. Hypoproliferative spheroids showed more migratory and no tumorigenic potential as compared to the hyperproliferative spheroids manifesting functional heterogeneity. Consistent with the functional characteristics in hypoproliferative spheroids, we observed the enrichment of EGR1 (cluster 0), IL1A (cluster 1), KRT6A (cluster 2), TPM1(cluster 5) and VIM (cluster 5), the genetic features mostly related to metastatic HNC (13–17). Notably, UMSCC-22B cell line is a hypopharyngeal squamous cell carcinoma derived from the metastatic lymph node, and this gene expression pattern reflects its EMT-like state. Overexpression of CCND2 (cluster 5), SMC4 (cluster 1), FABP5 (cluster 1), and NDRG1 (cluster 2 and 3) were observed, which are associated with proliferation in HNC (18–21). Interestingly, there was also overexpression of cell cycle inhibitor (CCND2: cluster4), kinase inhibitor (DUSP5: cluster 0), and tumor suppressor (S100A2: cluster 1), which might explain the lower proliferation rate in hypoproliferative spheroids (22–24). The dominance of fibroblast cells (highest CellKb hits ranks 1791), mesenchymal cells (highest CellKb hits ranks 457), and keratinizing barrier epithelial cells (CellKb hits ranks 1121.49) are associated with metastatic phenotype and observed in hypoproliferative spheroids (25,26). Hypoproliferative spheroids also contained genes related to stemness (KRT15 and ALDH3A1), consistent with stem cell-like characteristics, promoting the formation of spheroids. In hyperproliferative spheroid, clusters contained proliferative genes (for example, TOP2A, AURKA, CKS2, AREG, and ANLN), resembling a program of excessive proliferation in advanced HNC (27–31). Cell type landscape of all hyperproliferative spheroids clusters (0–3) was dominated by epithelial cells, neoplastic cells (highest CellKb hit rank 5515.99), and basal cells (highest CellKb hit rank 2081.23) aptly reflected its highly proliferative characteristics **(Fig. 7E and F).** We also observed several shared gene expressions of stem cell-like and proliferative properties across hypo- and hyperproliferative spheroids (KRT15, NDRG1, and SMC4).

Next, we investigated the potential biological functions associated with DEGs in different clusters across hypo-and hyperproliferative spheroids using GO enrichment analysis. The upregulated DEGs for hypoproliferative spheroids (hypermigratory) were specifically enriched in pathways such as signal transduction, cell migration, wound healing-related pathway, VEGFR, EGFR and TGFR beta signaling. While the hyperproliferative spheroids (hypomigratory) were enriched in pathways such as cell division, cell proliferation, EGFR, Wnt and Notch signaling, MAPK, ERK1 and ERK2 cascade **(Fig.7 G and H).** Cluster-specific analysis of DEGs showed that the hypoproliferative spheroids (cluster 5) were enriched with genes VIM, ODC1, and ID1 (associated with metastatic phenotype). Concomitant with that, cell type annotation predicted dominance of fibroblast cells and enrichment of pathways related to cell migration, TGF-beta, and VEGF signaling, suggesting an invasive phenotype. Our study also identified a KRT6A and KRT6B enriched epithelial cell state in cluster 2, while its cell type annotation revealed keratinizing barrier epithelial cells along with epithelial, basal, and neoplastic cells. Further, we mapped the top ten up- and down-regulated genes from scRNA seq data with The Cancer Genome Atlas Program (TCGA) data. We used a TNM plot to determine the cumulative expression of the top ten up- and down-regulated genes in each cluster of hypo- and hyperproliferative spheroids. We found that cluster 3 of hypoproliferative and cluster 1 of hyperproliferative spheroids cumulative expression of the top ten up- and down-regulated genes showed significant up and down-regulation in tumor samples (n = 29,376) as compared to adjacent normal tissue (n = 3691) (32) (**Fig. 7I** and **J**, **Supplementary Fig. 10 and 11**). Surprisingly, we found that cluster 3 of hypoproliferative spheroids were uniquely enriched in pathways, such as cell migration, ERK1 and ERK2 cascade.

Our scRNAseq results suggested that clusters 2 and 5, in addition to cluster 3 as suggested by TCGA analysis might be responsible for hypermigration in hypoproliferative spheroids. Similarly, in hyperproliferative spheroid, cluster 1 showed marker genes for cell proliferation, such as TOP2A, AURKA, CKS2, HMGB2, and SMC4, which correlated positively with the cell type annotation of neoplastic cells with the enrichment of EGFR, MAPK and Wnt signaling pathways. Thus, our scRNAseq and TCGA findings suggest that cluster 1 might be involved in the fast growth behaviour of the hyperproliferative spheroids.

## Discussion

Fundamental characteristics of human malignancies, such as heterogeneity molecular diversity and cellular plasticity, are crucial to cancer development. Considerable efforts have been made to understand the high level of heterogeneity in HNC at the single-cell level and its role in therapeutic resistance (33–37). Although established cell lines are frequently utilized in cancer research, it is unclear whether they accurately represent the cellular heterogeneity observed in the tumors. We attempted to describe the landscape of cellular variability within HNC cell lines to investigate the heterogeneity within the various cancer cells and their capacity to replicate intratumoral heterogeneity in patients with HNC.

Here, we developed single-cell derived clonal spheroids from model cell lines in HNC. These spheroid models were developed to identify the cellular diversity that can drive and support tumorigenesis in 3D and enable the molecular interrogation of structural and functional heterogeneity in HNC tumors. The diverse growth kinetics of the sub-clonal population within the cell line enabled us to sort the spheroids into hypoproliferative (smaller diameter) and hyperproliferative (larger diameter) spheroids. We demonstrated a similar trend of structural variability and relevant molecular patterns in spheroid formation from the single-cells isolated from the tumors of HNC patients. Recent research in the SCDS has identified similar structural variations and has reinforced the occurrence of structural heterogeneity within these clonal spheroids (5, 38).

Interestingly, we observed variability in the frequency of spheroid formation at different stages of cancer progression with varying samples of patients. We considered UMSCC-22B cells as late-stage tumor cells as it was derived from lymph node metastasis. Interestingly, in cells isolated from all late-stage tumors (T3 and T4), the frequency of spheroid formation was higher for a hypoproliferative spheroid as compared to a hyperproliferative spheroid. However, in comparatively early-stage tumor (T2), vice-versa was observed. Similar studies with higher sample sizes may better estimate the frequency of such spheroids. Further, structural-functional studies revealed variation in migratory and tumorigenic potential of single-cell derived hypo-and hyperproliferative spheroids. Hypoproliferative spheroids showed more migratory potential but could not initiate tumor growth in mice whereas the hyperproliferative spheroids had tumorigenic potential but lesser migratory potential.

Consequently, we observed an increased fluorescence intensity for the percentage of dead cells in hyperproliferative spheroids derived from patient cells after staining with EtBr in comparison with hypoproliferative spheroids after treatment with 4 Gy, Cisplatin, and Cisplatin along with 4 Gy radiation. This may be attributed to the fact that hyperproliferative spheroids are comprised of rapidly proliferating cells compared to hypoproliferative spheroids. It has been previously reported that rapidly proliferating cells are more sensitive to radiation, leading to an enhanced DNA damage response (39). Also, the fact that quiescent cells are known to be more resistant to radiation and chemotherapy than cells growing in a more proliferative environment may predict the resistance. Earlier studies reported that quiescent cells can undergo better repair of potentially lethal damage concerning proliferating cells, and these mechanisms could be responsible for the observed differences after treatment (40).

Strategies to identify molecular players in the complex structural-functional basis of heterogeneity among single-cell derived hypo-and hyperproliferative spheroids might be an important strategy towards understanding tumor development in HNC. However, very limited studies were done to justify spheroids as a preclinical research model for HNC at a large-scale transcriptomic level. Our single-cell landscape of SCDS addresses this lacune and identifies many prospective targets for the prevention, detection and treatment. The cellular process involving VEGFR, EGFR, TGFR beta, Wnt, Notch, and MAPK signaling cascade with their complex gene interaction network was the critical contributors in the functional heterogeneity observed in hypo-and hyperproliferative spheroids.

This study further revealed clinically relevant sub-populations in hypopharyngeal carcinoma along with their molecular indicators. Analysis of both the clusters and DEGs revealed the transcriptional features affecting the structure-functional variations in hypo- and hyperproliferative spheroids. We identified metastatic molecular signature in hypoproliferative spheroids; however, it was also accompanied by markers associated with kinase inhibitors, tumor suppressors, and cell cycle inhibitors, explaining its high migratory potential and slower growth rate. Variations in cell type composition within the clusters of both hypo- and hyperproliferative spheroids implicate the complex interactions within the tumor microenvironment driving cancer progression. These may be distinct sub-populations of cells or a continuous spectrum of cellular states that evolved during the clonal expansion of the tumor. These hypo-and hyperproliferative spheroids had intact transcriptional memory of the primary tumor at the hypopharynx. Cell type analysis indicated the presence of keratinizing barrier epithelium cells, and corresponding to this cell type, various keratins were enriched in these clusters, which is reminiscent of the stratified epithelium of the mucosal lining in the upper aerodigestive tract (1). Squamous cell carcinoma, which accounts for more than 90% of the cancers of the head and neck, arises from the squamous dysplasia at the surface epithelium of these regions (41). Further studies are required to elucidate and verify the structural and functional characteristics of these single-cell-derived clonal spheroids, which is an emerging preclinical model for HNC to study tumor heterogeneity.

In conclusion, extensive variability in gene expression was identified between hypo-and hyperproliferative spheroids, manifesting tumor heterogeneity in the HNC. Cancer biology research is increasingly focusing on 3D spheroid culture. In this study, we have developed single-cell-derived clonally propagated spheroids, which could aid in understanding the rare cell population responsible for tumor maintenance and drug resistance. The significant advantages of a SCDS are that we can see how cells grow, proliferate, interact, and absorb nutrients and chemical compounds at the single-cell level. The SCDS could be used as a tool to develop the next generation of therapeutics, determine the optimal therapeutics, and predict the progression of cancer.

## Authors’ contributions

**J. Pandey:** Conceptualization, data curation, software and formal analysis, investigation, visualization, methodology, writing–original draft, writing-review, and editing. **M. Malik:** Conceptualization, analysis and interpretation of data including statistics, biostatistics, and computational analysis, visualization, writing review, and editing. **R. Shyanti:** Conceptualization, data curation, software and formal analysis, investigation, visualization, methodology, writing–original draft. **P. Parashar:** Conceptualization, data curation, software and formal analysis, investigation, visualization, methodology, writing and editing. **P. Kujur:** Formal analysis, investigation, visualization, writing–original draft. **D. Mishra:** Data curation, investigation, visualization, methodology, writing and editing. **D. Tailor:** Formal analysis and investigation, writing and editing. **J. Lee:** Formal analysis and investigation, writing and editing. **T. Kataria:** Data curation, formal analysis, managed patients, investigation, and resources. **D. Gupta:** Data curation, formal analysis, and investigation, managed patients and resources. **H. Verma:** Data curation, formal analysis, and investigation, managed patients, and resources. **S. Malhotra:** Conceptualization, resources, supervision, funding acquisition, writing and editing. **S. Kateriya:** Formal analysis, visualization and editing. **V. Tandon:** Conceptualization, formal analysis, funding acquisition, study supervision, project administration, writing and editing. **R. Chaturvedi:** Conceptualization, formal analysis, funding acquisition, study supervision, project administration, writing and editing. **R. Singh:** Conceptualization, formal analysis, funding acquisition, study supervision, project administration, writing and editing.

## Acknowledgments

The work was supported by DPRP-Department of Science & Techhnology (VI-D&P/546/2016-17/TDT (C)), Indo-US Science Technology Forum (IUSSTF/JC-131/2019), and Indian Council of Medical Research (F.No. 6/11/23/dr/icmr), India. The in part support is also acknowledged from DBT-Builder, DST-PURSE and UGC-DRS programs. We thank All India Institute of Medical Sciences, New Delhi, Division of Radiation Oncology, Medanta-The Medicity, Gurgaon, India for providing research facilities and patient samples, and Jawaharlal Nehru University, New Delhi for Central Laboratory Animal Facility. Jyoti Pandey, Palak Prashar, and Deepali Mishra were supported by CSIR-UGC-JRF, ICMR-SRF, and UGC-JRF, respectively, India. We would like to thank Navneendra Singh for patient sample collection and animal breeding. The funders of the study had no role in study design, data collection, analysis, interpretation, or writing of the manuscript. V. Tandon, R. Chaturvedi, R. Singh had full access to all data in the study and as corresponding authors had final responsibility for the decision to submit for publication.

## Institutional Review Board Statement

All animal experiments adhered to institutional protocols and were conducted within the specific pathogen-free facility CLAR at the Jawaharlal Nehru University, India, following an approved Institutional Animal Ethics Committee (IAEC) denoted as (IAEC Code no. 05/2017). Head and neck squamous cell carcinoma (HNSCC) tumor samples were collected from Indian patients at All India Institute of Medical Sciences (AIIMS), New Delhi, following the IRB approval (No. 2017-123) and also from Medanta Hospital, Gurugram, India following its IRB approval (No. MICR-1013/2019). Certified pathologists interpreted the histology of the specimens, and the tumor staging was determined according to the American Joint Committee on Cancer (AJCC) staging manual, 7th edition.

