## Supplementary files for "Clonal spheroids capture functional and genetic heterogeneity of head and neck cancer"

**Supplementary figures:**

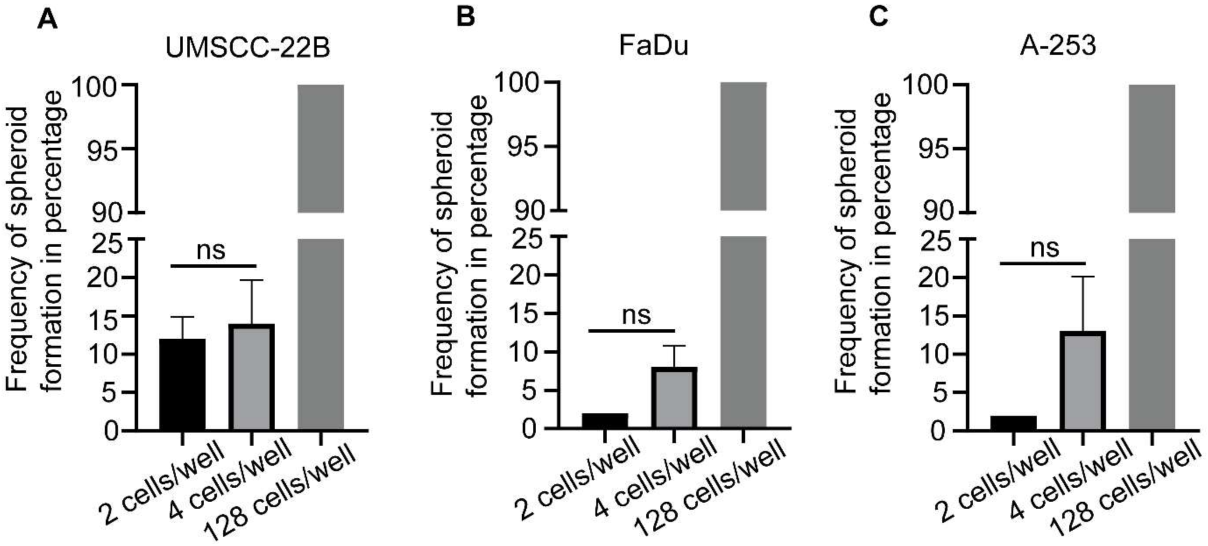


**Supplementary Figure 1.** The **frequency of spheroid formation in ULA 96-well round bottom microplate increases with seeding density.** Spheroid forming frequency in **(A)** UMSCC-22B **(B)** FaDu **(C)** A-253 increased with the increase in seeding density (2, 4 and 128 cells/well). The data are represented as Mean ± SEM of three independent experiments. (ns, non-significant).

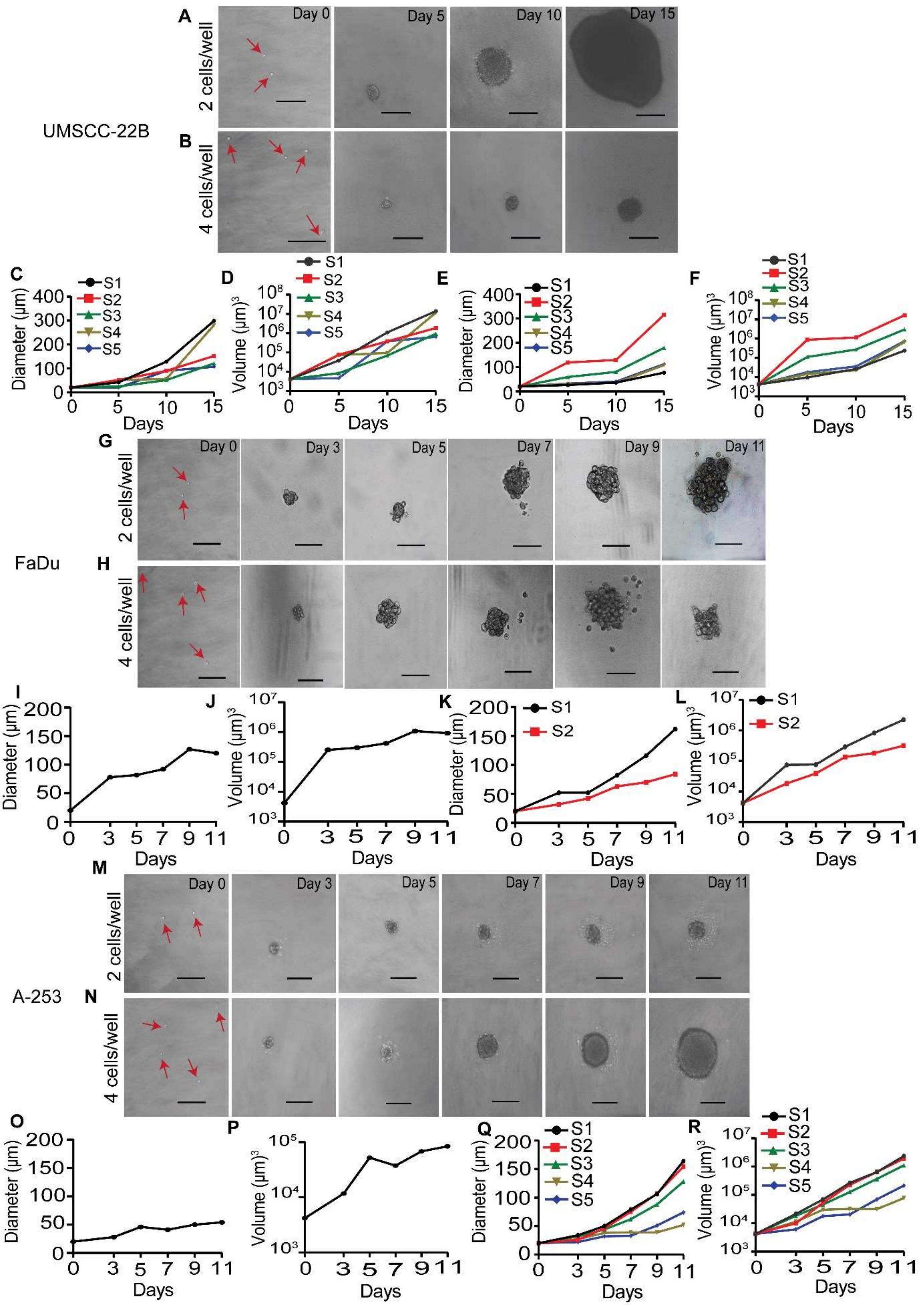


**Supplementary Figure 2. Morphology and growth kinetics of spheroids derived from 2 and 4 cells/well in the ULA round bottom plate**. **(A and B)** Representative microscopic images of UMSCC-22B spheroids captured at days 0, 5, 10, and 15. **(C-F)** Growth kinetics of UMSCC-22B spheroids in terms of diameter (X and Y planes) and volume (X, Y, and Z planes) (S1-5, Spheroid 1-5). **(G and H)** Representative microscopic images of FaDu spheroids were captured on days 0, 3, 5, 7, 9, and 11. **(I-L)** Growth kinetics of FaDu spheroids in terms of diameter (X and Y planes) and volume (X, Y, and Z planes) (S1-2, Spheroid 1-2).. **(M and N)** Representative microscopic images of A-253 spheroids captured at days 0, 3, 5, 7, 9, and 11. **(O-R)** Growth kinetics of A-253 spheroids in terms of diameter (X and Y planes) and volume (X, Y, and Z planes) (S1-5, Spheroid 1-5).. (10X objective, scale bar = 100 μm).

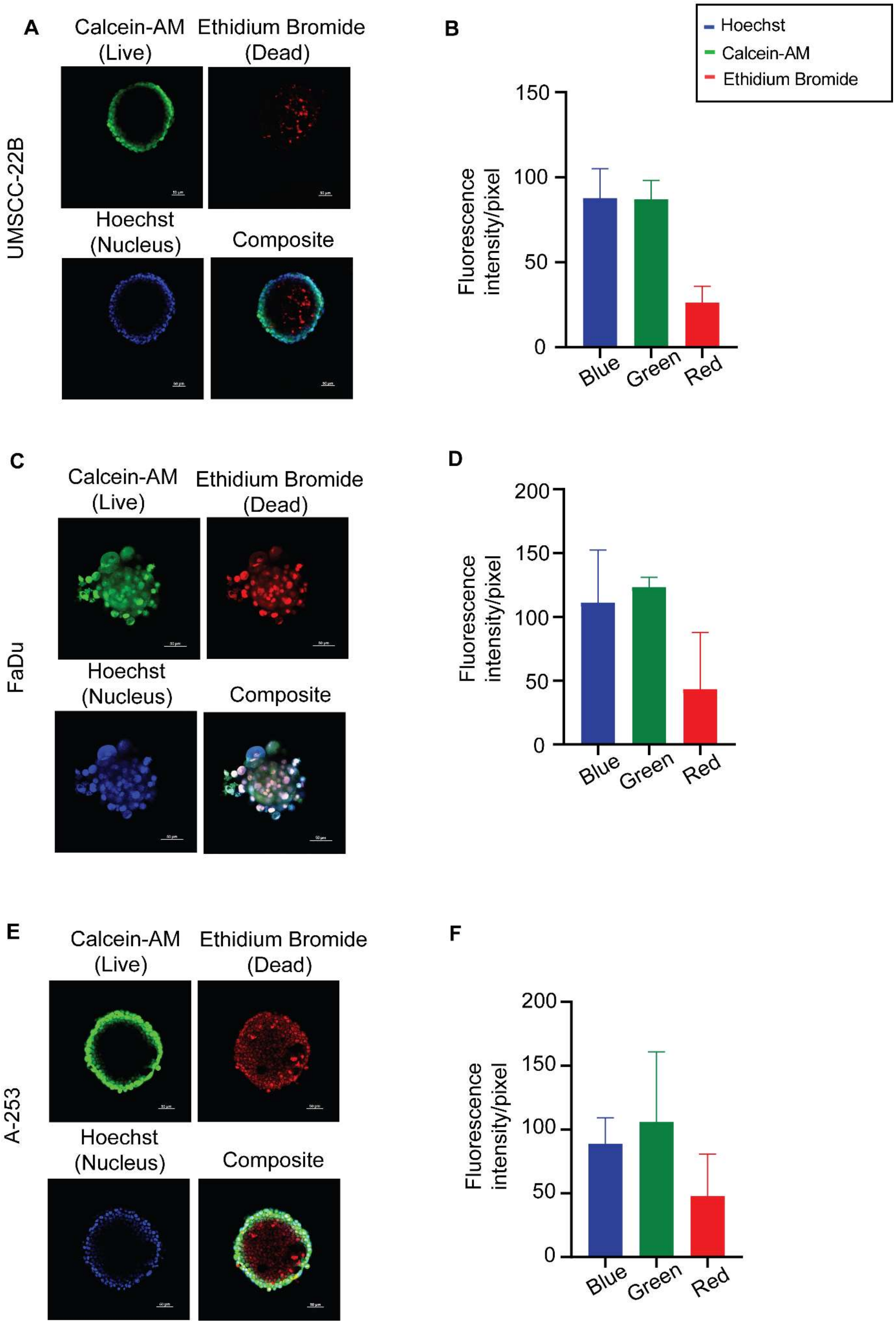
A
Calcein-AM (Live)
Ethidium Bromide (Dead)
UMSCC-22B
Hoechst (Nucleus)
Composite
B
Hoechst
Calcein-AM
Ethidium Bromide
Fluorescence intensity/pixel
150
100
50
0
Blue
Green
Red
C
Calcein-AM (Live)
Ethidium Bromide (Dead)
FaDu
Hoechst (Nucleus)
Composite
D
Fluorescence intensity/pixel
200
150
100
50
0
Blue
Green
Red
E
Calcein-AM (Live)
Ethidium Bromide (Dead)
A-253
Hoechst (Nucleus)
Composite
F
Fluorescence intensity/pixel
200
150
100
50
0
Blue
Green
Red

**Supplementary Figure 3. Confocal imaging of spheroid derived from 4 cells/well in ULA 96-well round bottom microplate displays typical spheroid morphology**. Representative confocal images of (**A)** UMSCC-22B **(C)** FaDu **(E)** A-253 cells derived spheroid. Calcein-AM (live, green), Ethidium Bromide (dead, red), and Hoechst-33342 (nuclear, blue) were used for staining. **(B, D and E)** Quantification of fluorescence intensity of dyes staining (**B)** UMSCC-22B **(D)** FaDu **(F)** A-253 spheroids and fluorescence intensity/pixel is represented as the mean ± SEM obtained from n=3 spheroids. 10X and 20X magnification, scale bar = 50 μm.

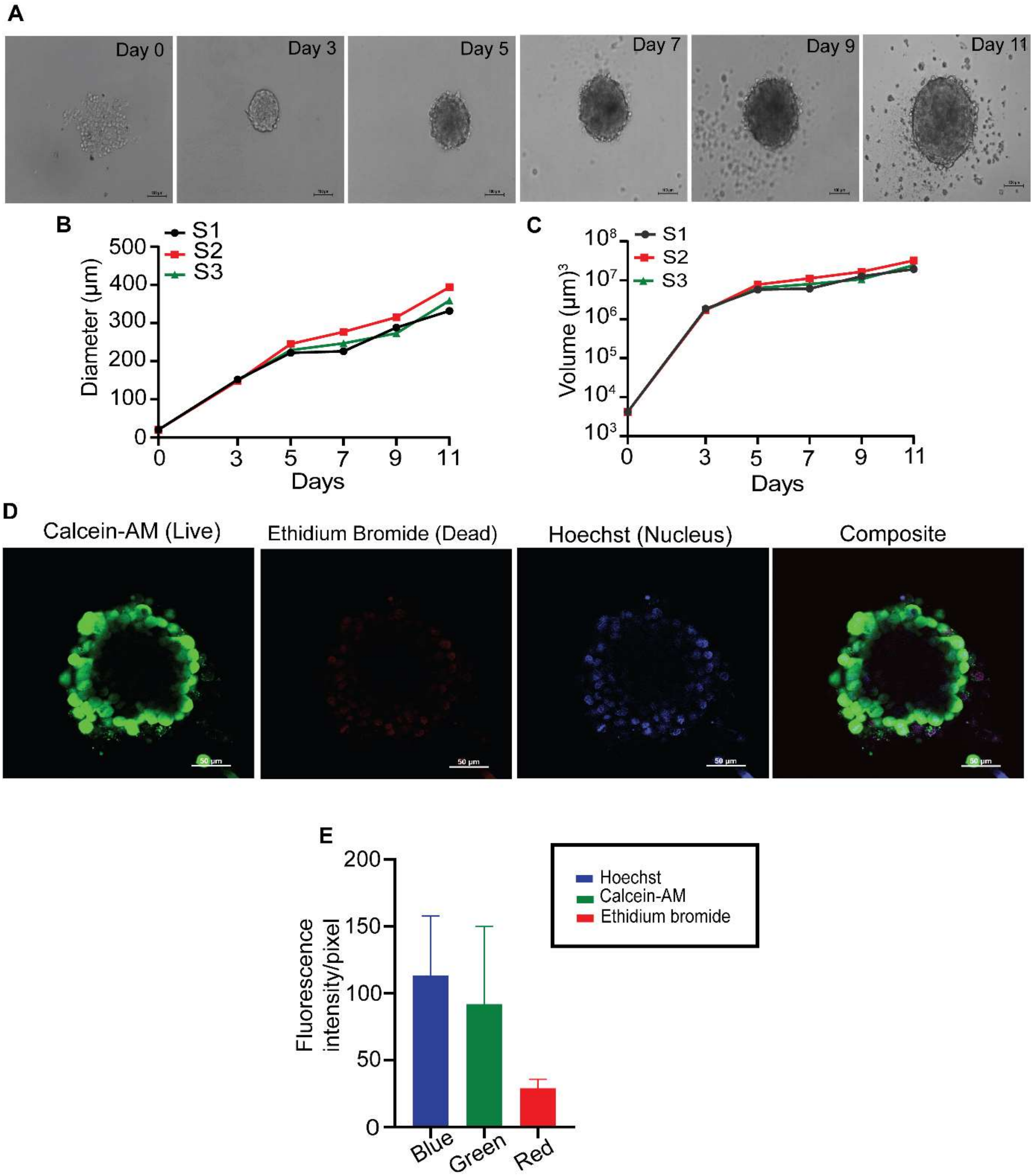


**Supplementary Figure 4. Morphology and growth kinetics of FaDu spheroids derived from 128 cells/well. (A)** Representative spheroids images were capturedon days 0, 3, 5, 7, 9, and 11 (10x magnification, scale bar = 100 μm). (**B and C**) Growth kinetics of spheroids in terms of diameter (X and Y planes) and volume of spheroids (X, Y, and Z planes) (S1-3, Spheroid 1-3). (**D**) Representative confocal images of FaDu cells derived spheroid. Calcein-AM (live, green), Ethidium Bromide (dead, red), and Hoechst-33342 (nuclear, blue) were used for staining. **(E)** Quantification of fluorescence intensity of dyes staining FaDu derived spheroid and fluorescence intensity/pixel is represented as the mean ± SEM obtained from n=3 spheroids. 20 X magnification, scale bar = 50 μm.

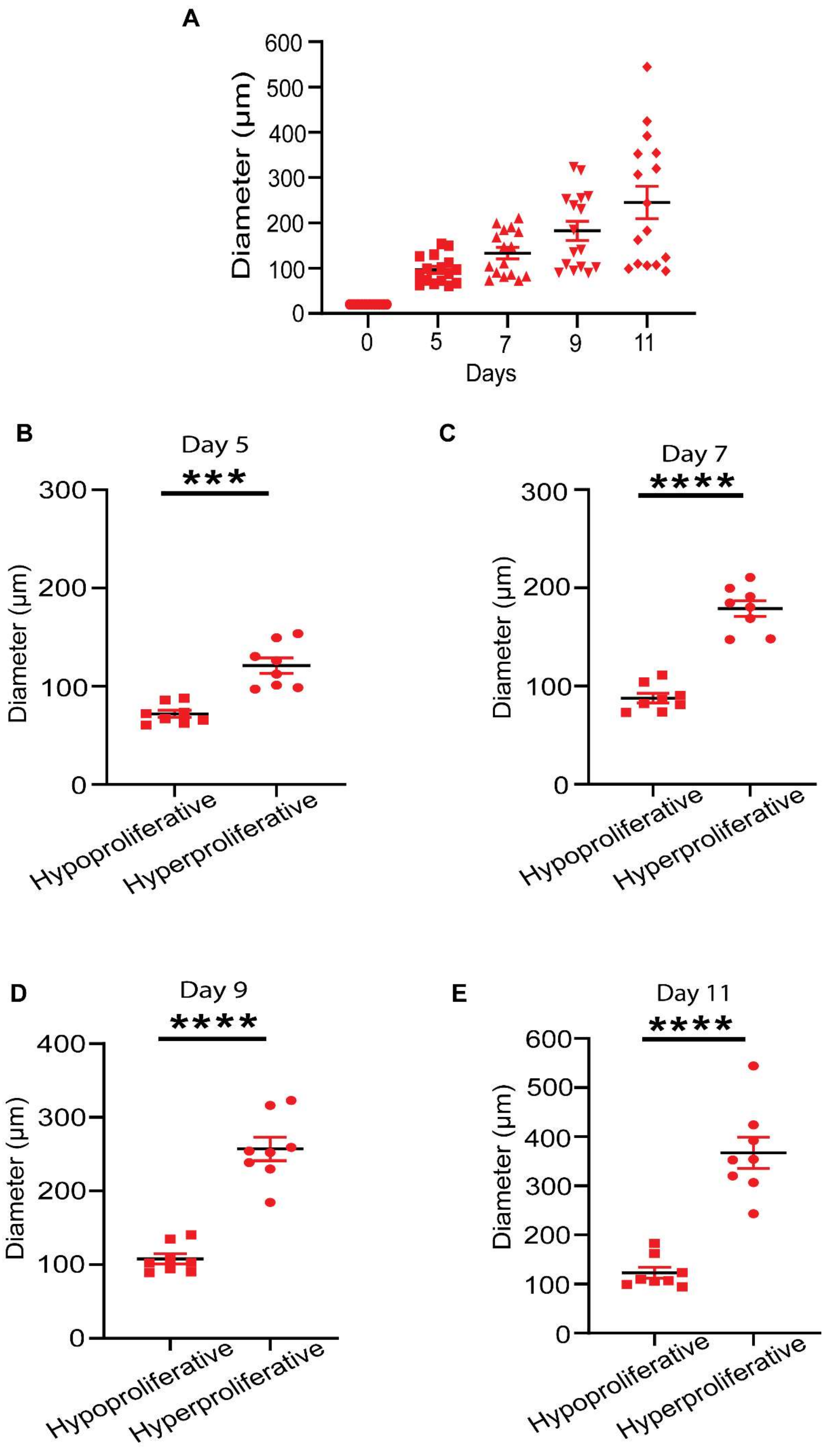

A
Diameter (µm)
600
500
400
300
200
100
0
0
5
7
9
11
Days
B
Day 5
***
300
200
100
0
Diameter (µm)
Hypoproliferative
Hyperproliferative
C
Day 7
****
300
200
100
0
Diameter (µm)
Hypoproliferative
Hyperproliferative
D
Day 9
****
400
300
200
100
0
Diameter (µm)
Hypoproliferative
Hyperproliferative
E
Day 11
****
600
500
400
300
200
100
0
Diameter (µm)
Hypoproliferative
Hyperproliferative

**Supplementary Figure 5.** Scatter plot showing the diameter of SCDS. **(A)** Combined diameter of hypo-and hyperproliferative spheroids. **(B)** Diameter of hypo- and hyperproliferative on day 5. **(C)** on day 7. **(D)** on day 9. **(E)** on day 11. The error bar represents SEM (n=8). The statistical method employed was a paired t-test.

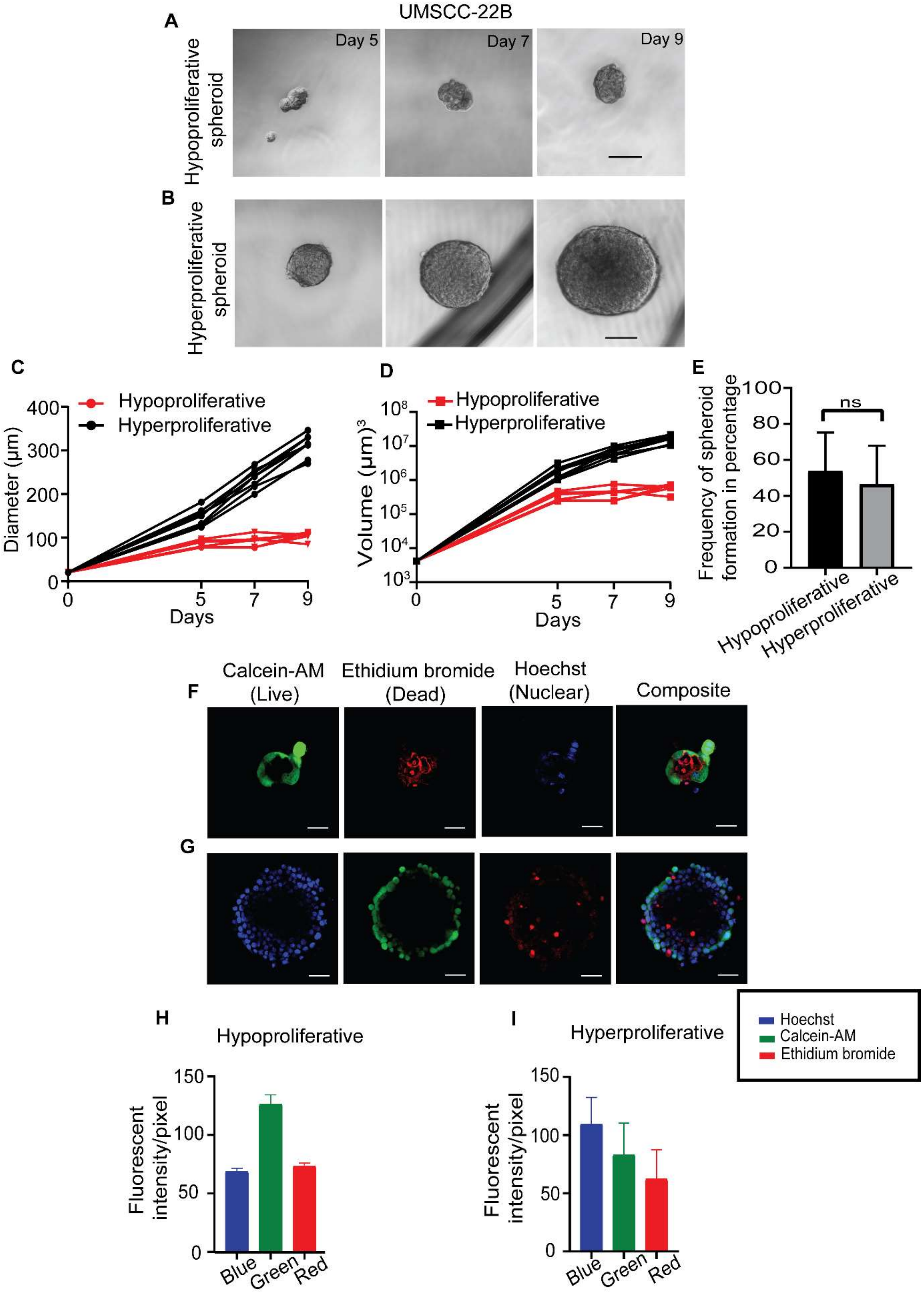


**Supplementary Figure 6. Morphology and growth kinetics of UMSCC-22B derived hypo- and hyperproliferative spheroids in a ULA 96-well flat bottom plate. (A and B)**

Representative images of UMSSC-22B hypo-and hyperproliferative spheroids were captured on days 5, 7, and 9 (10X magnification, scale bar = 100 µm). (**C and D**) Growth kinetics of hypo-and hyperproliferative spheroids in terms of diameter (X and Y planes) and volume (X, Y, and Z planes). **(E)** Frequency of spheroids formation of hypo-and hyperproliferative spheroids. **(F and G)** Representative confocal images of FaDu cells derived hypo-and hyperproliferative spheroids. Calcein-AM (live, green), Ethidium Bromide (dead, red), and Hoechst-33342 (nuclear, blue) were used for staining. **(E)** Quantification of fluorescence intensity of dyes staining UMSCC-22B cells derived hypo-and hyperproliferative spheroids and fluorescence intensity/pixel is represented as the mean ± SEM obtained from n=3 spheroids. 20x magnification, scale bar = 50 µm.

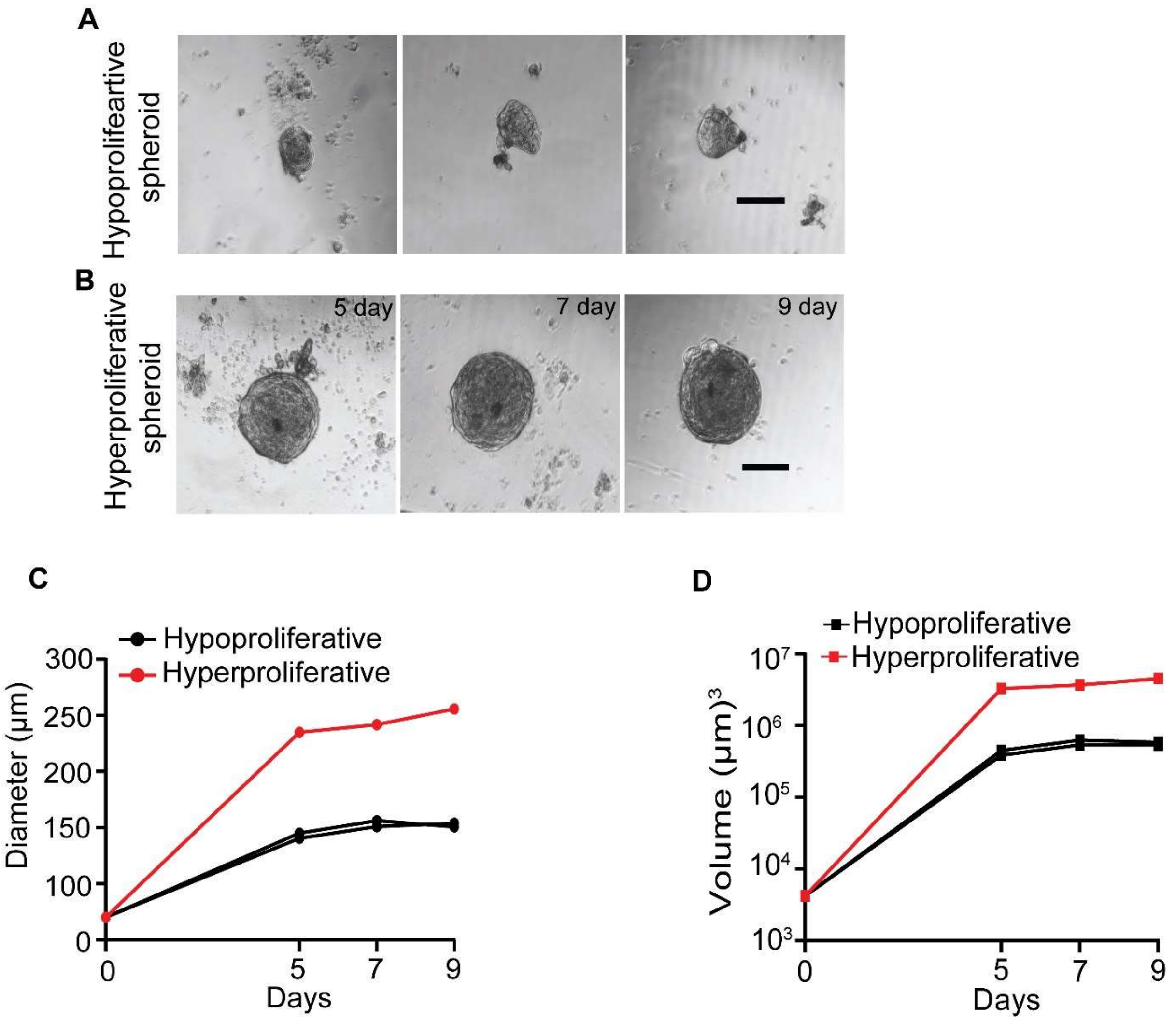


**Supplementary Figure 7. Generation of hypo-and hyperproliferative spheroids using HNC patient single-cell suspension in a ULA flat bottom plate. (A and B)** Representative microscopic images of hypo-and hyperproliferative spheroids at days 5, 7 and 9. **(C and D)**

Growth kinetics of hypo-and hyperproliferative spheroids in terms of diameter (X and Y planes) and volume (X, Y, and Z planes).

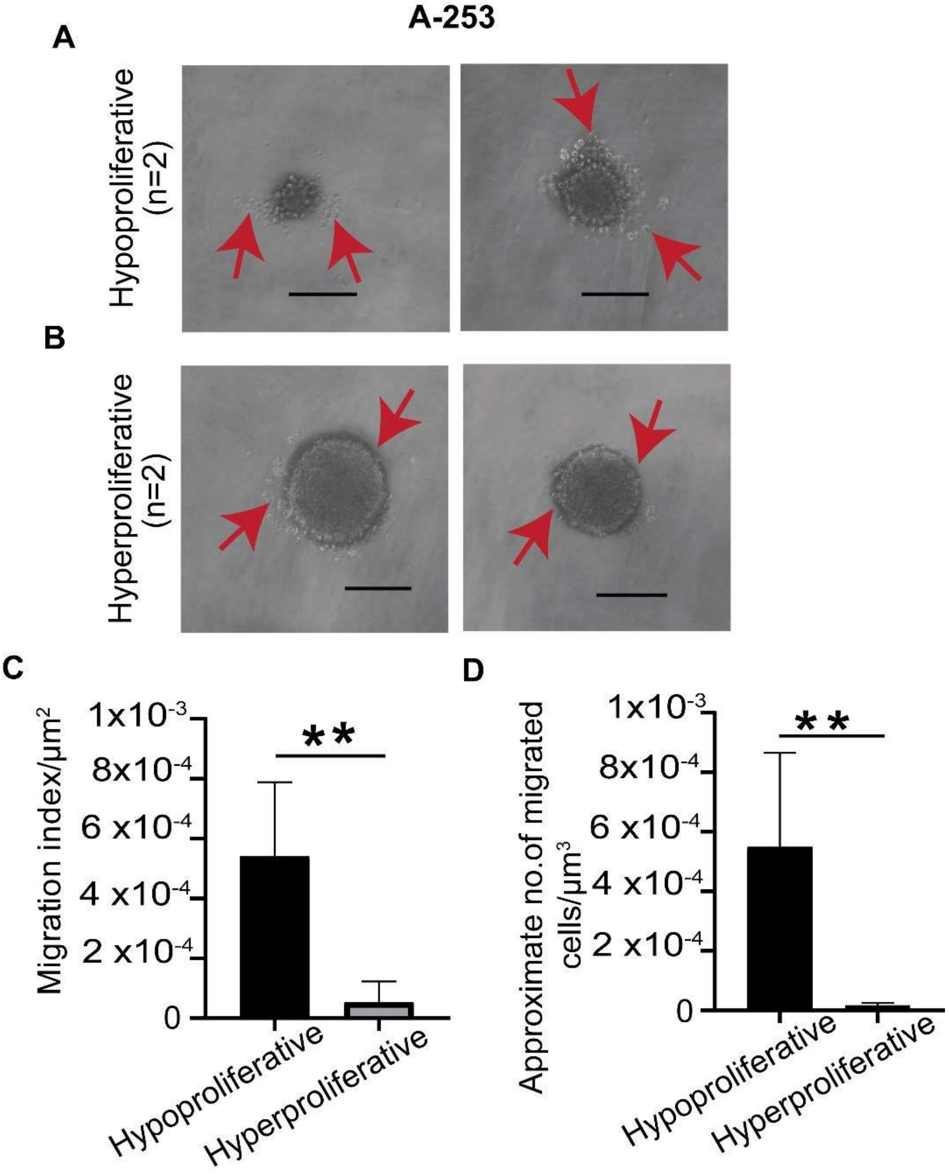


**Supplementary Figure 8. 3D single-cell spheroid migration assay reveals functional heterogeneity in hypo-and hyperproliferative spheroids generated from A-253 cells. (A and B)** Representative images of cell migration from hypo-and hyperproliferative spheroids (10X magnification, scale bar = 100 µm). **(C)** Quantification of migration using migration index of hyperproliferative and hypoproliferative spheroid. **(D)** Quantification of no. of migrated cells from the periphery of spheroids. (ns-non-significant, $^{**}p \leq 0.01$).

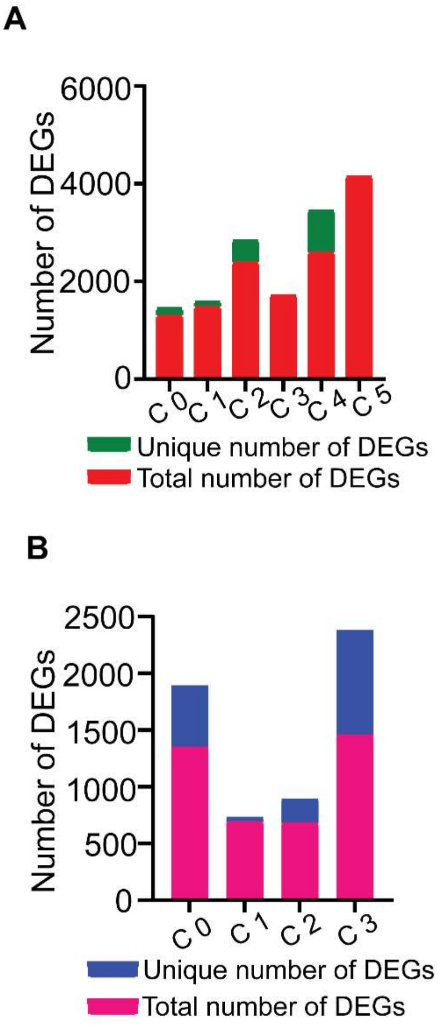


**Supplementary Figure 9. (A and B)** The bar plots depict the total and unique DEGs of each identified cluster in hypo-and hyperproliferative spheroid.

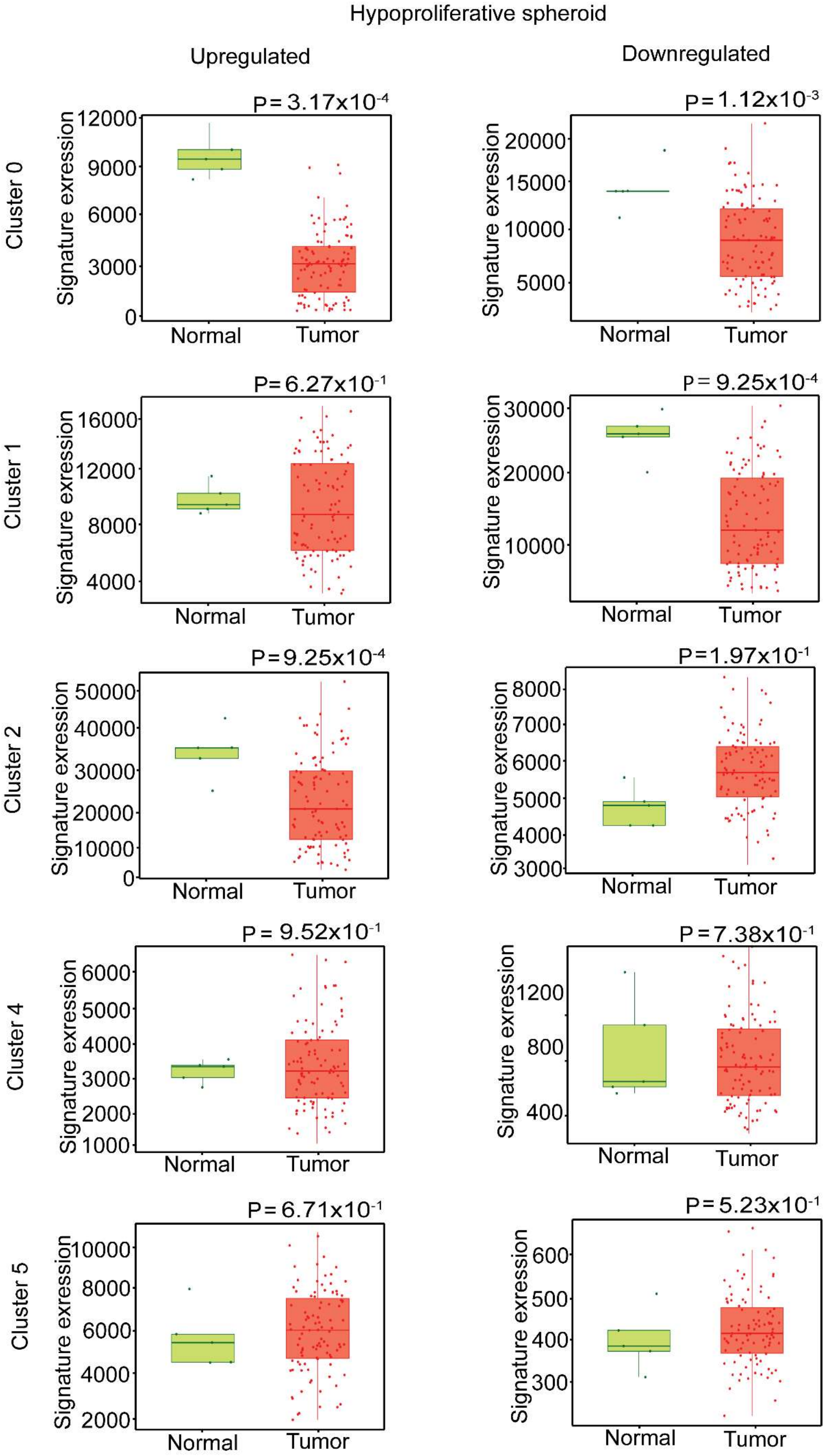

Hypoproliferative spheroid
Upregulated
Downregulated
Cluster 0
Cluster 1
Cluster 2
Cluster 4
Cluster 5
P=3.17x10-4
P=1.12x10-3
P=6.27x10-1
P=9.25x10-4
P=9.25x10-4
P=1.97x10-1
P=9.52x10-1
P=7.38x10-1
P=6.71x10-1
P=5.23x10-1
Signature exression
Normal
Tumor

**Supplementary Figure 10. (A and B)** Bar graph depicting the up and downregulated genes in different clusters (clusters 0, 1, 2, 4, and 5) of hypoproliferative spheroid.

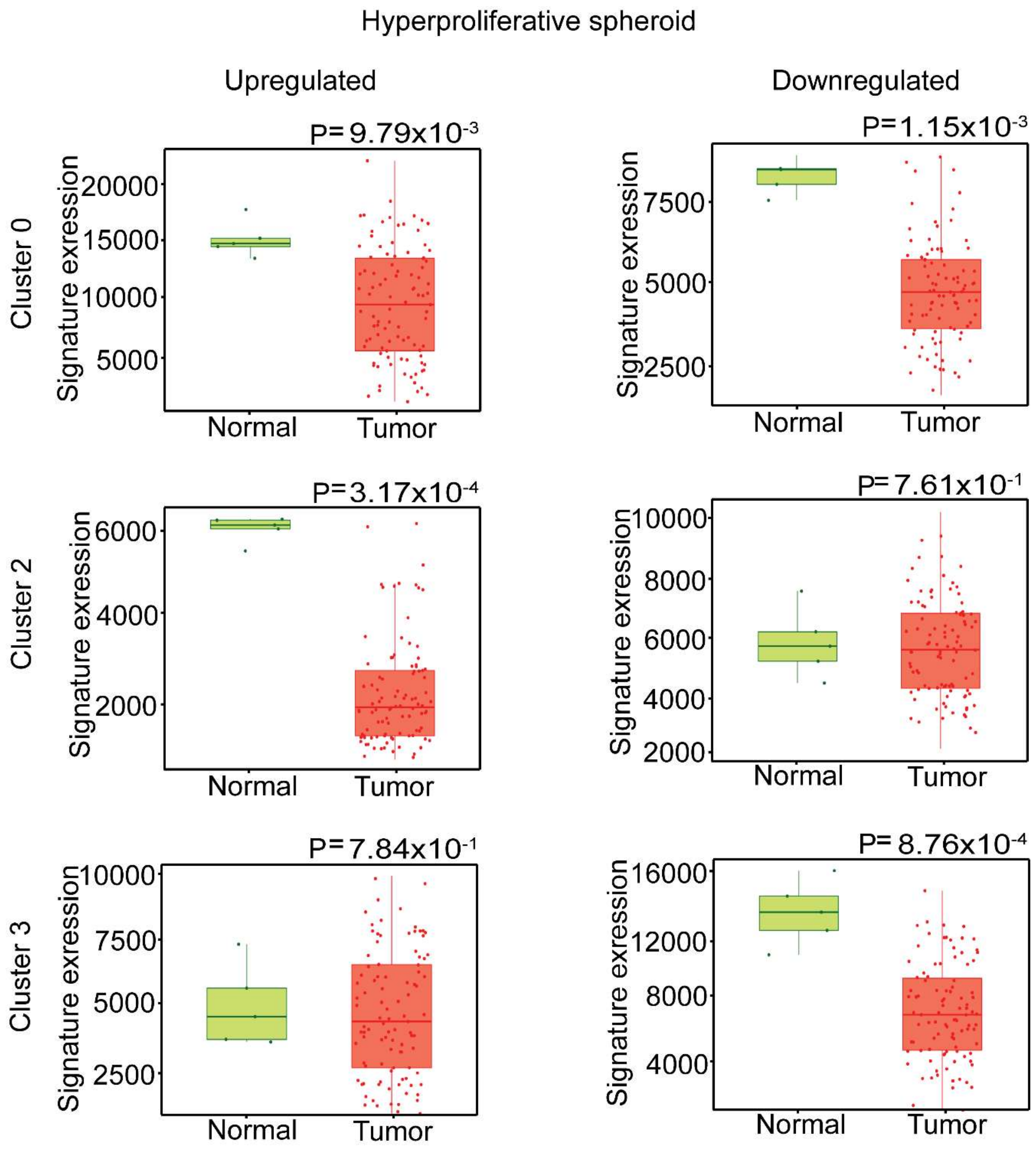


**Supplementary Figure 11. (A and B)** Bar graph depicting the up and downregulated genes in different clusters (clusters 0, 2, and 3) of the hyperproliferative spheroid.
