## Supplementary tables for "Clonal spheroids capture functional and genetic heterogeneity of head and neck cancer"

**Supplementary table 1: Total and unique no. of DEGs present in each cluster of hypoproliferative spheroids**

Related to figure 7A

| Clusters | Total no. of DEGs | Unique no. of DEGs |
|---|---|---|
| **0** | 1314 | 150 |
| **1** | 1492 | 105 |
| **2** | 2398 | 449 |
| **3** | 1726 | 0 |
| **4** | 2604 | 851 |
| **5** | 4155 | 0 |

**Table 2:** Total and unique no. of DEGs present in each cluster of hyperproliferative spheroids

Related to figure 7B

| Clusters | Total no. of DEGs | Unique no. of DEGs |
|---|---|---|
| **0** | 1349 | 540 |
| **1** | 693 | 38 |
| **2** | 683 | 207 |
| **3** | 1455 | 921 |

**Table 3:** Differentially expressed genes of each cluster of hypoproliferative spheroid

Related to Figure 7C

| (A) | Cluster 0 | |
|---|---|---|
| **Gene** | ***P* value** | **Average log2FC** |
| **KRT13** | 6.20E-113 | 2.481635894 |
| **MT-CO2** | 4.85E-47 | 1.057118 |
| **MT-CO1** | 2.47E-53 | 1.151088762 |
| **MT-CO3** | 1.94E-51 | 1.134036492 |
| **MDK** | 1.05E-68 | 1.206016765 |
| **KRT15** | 4.49E-67 | 1.563960459 |
| **ALDH3A1** | 2.42E-37 | 1.261246822 |
| **EGR1** | 2.59E-31 | 1.32541404 |
| **DUSP5** | 5.31E-32 | 1.154298669 |
| **MT-ATP6** | 2.64E-50 | 1.162043576 |

| (B) | Cluster 1 | |
|---|---|---|
| **Gene** | ***P* value** | **Average log2FC** |
| **CAV1** | 2.51E-82 | 1.755459 |
| **EIF1AX** | 9.60E-53 | 0.947118 |
| **FABP5** | 1.67E-45 | 0.928984 |
| **IL1A** | 2.40E-45 | 1.346947 |
| **TUBA1B** | 7.02E-44 | 1.080077 |
| **CCL20** | 3.34E-35 | 1.392613 |
| **S100A2** | 2.16E-34 | 0.92984 |
| **SMC4** | 1.19E-35 | 0.99772 |
| **MT2A** | 7.13E-34 | 1.259861 |
| **HIST1H4C** | 3.73E-23 | 0.979365 |

| (C) | Cluster 2 | |
|---|---|---|
| **Gene** | ***P* value** | **Average log2FC** |
| **SPRR1B** | 2.13E-81 | 5.097446 |
| **S100A8** | 1.381E-103 | 4.931207 |
| **S100A9** | 4.90E-87 | 3.424656 |
| **PI3** | 2.973E-114 | 4.834687 |
| **POLR2J3** | 3.28E-94 | 3.031692 |
| **S100A7** | 3.18E-43 | 3.016662 |
| **KRT6A** | 5.08E-62 | 3.978481 |
| **KRT6B** | 3.74E-41 | 3.179217 |
| **P4HA1** | 6.40E-10 | -0.90784 |
| **NDRG1** | 1.80E-24 | 1.07161 |

| (D) | Cluster 3 | |
|---|---|---|
| **Gene** | ***P* value** | **Average log2FC** |
| **PLOD2** | 1.55E-52 | 1.774588 |
| **CASP14** | 1.50E-25 | 1.896099 |
| **SCD** | 1.22E-28 | 1.497395 |
| **UCA1** | 1.05E-20 | 1.457285 |
| **SLC5A3** | 5.49E-30 | 1.523327 |
| **CALB1** | 2.76E-12 | 1.769368 |
| **DNAJB9** | 4.28E-37 | 1.389893 |
| **AGR2** | 1.56E-13 | 1.383642 |
| **P4HA1** | 2.45E-64 | 1.90771 |
| **NDRG1** | 5.97E-44 | 1.91007 |

| (E) | Cluster 4 | |
|---|---|---|
| **Gene** | ***P* value** | **Average log2FC** |
| **UCHL1** | 1.073E-103 | 3.303057 |
| **SLC25A21** | 7.41E-89 | 2.205439 |
| **MGST1** | 2.82E-99 | 2.496415 |
| **CDKN2A** | 8.433E-122 | 2.351122 |
| **TRNP1** | 5.57E-83 | 2.206427 |
| **SNHG25** | 3.85E-92 | 2.135215 |
| **CKB** | 4.21E-82 | 2.084907 |
| **GLS** | 2.59E-90 | 1.956704 |
| **NQO1** | 5.54E-84 | 1.940341 |
| **FTL** | 1.09E-70 | 1.872465 |

| (F) | Cluster 5 | |
|---|---|---|
| **Gene** | ***P* value** | **Average log2FC** |
| **VIM** | 1.06E-74 | 4.900315413 |
| **ODC1** | 1.044E-124 | 3.733199188 |
| **UBB** | 2.40E-76 | 3.207936185 |
| **PMP22** | 1.36E-71 | 3.276472412 |
| **CCND2** | 3.29E-75 | 3.501590893 |
| **ZNF90** | 2.88E-121 | 3.514482987 |
| **TPM1** | 3.195E-128 | 3.555675064 |
| **SERPINE1** | 9.60E-66 | 4.145534186 |
| **EIF4A1** | 9.248E-131 | 3.037430947 |
| **ID3** | 6.92E-68 | 3.096989644 |

### Table 4: Differentially expressed genes of each cluster of hyperproliferative spheroid

Related to Figure 7D

| (A) | Cluster 0 | |
|---|---|---|
| **Gene** | ***P* value** | **Average log2FC** |
| **NDRG1** | 3.34E-33 | 1.795103 |
| **ERO1A** | 6.46E-23 | 1.505011 |
| **P4HA1** | 2.52E-21 | 1.188855 |
| **CP** | 3.76E-15 | 1.387663 |
| **LCN2** | 5.09E-16 | 1.297841 |
| **C3** | 1.33E-14 | 1.250857 |
| **CASP14** | 3.56E-12 | 1.349552 |
| **CALB1** | 2.40E-10 | 2.133158 |
| **PI3** | 5.97E-06 | 2.038952 |
| **KRT6A** | 0.002804 | 1.813816 |

| (B) | Cluster 1 | |
|---|---|---|
| **Gene** | ***P* value** | **Average log2FC** |
| **TUBA1B** | 6.41E-31 | 1.354825 |
| **SMC4** | 7.15E-29 | 1.253732 |
| **HMGB2** | 1.36E-31 | 1.471308 |
| **ANLN** | 7.43E-24 | 1.239846 |
| **CKS2** | 3.29E-26 | 1.173884 |
| **TPX2** | 4.67E-21 | 1.330709 |
| **TOP2A** | 2.19E-20 | 1.515042 |
| **AURKA** | 1.69E-19 | 1.308954 |
| **HIST1H4C** | 2.17E-16 | 1.303604 |
| **MT2A** | 1.30E-05 | 0.917532 |

| (C) | Cluster 2 | |
|---|---|---|
| **Gene** | ***P* value** | **Average log2FC** |
| **KRT13** | 6.61E-30 | 2.604844185 |
| **TNFSF10** | 2.80E-17 | 1.436137368 |
| **B4GALT5** | 4.99E-24 | 1.772594728 |
| **ID1** | 2.21E-15 | 1.525296919 |
| **SAMD9** | 1.46E-17 | 1.594590112 |
| **HES1** | 5.84E-12 | 1.366306874 |
| **SCGB1A1** | 2.43E-07 | 1.591301109 |
| **IFIT2** | 7.13E-05 | 1.641281811 |
| **KRT15** | 7.14E-16 | 1.331744459 |
| **IFIT3** | 1.19E-08 | 2.034699963 |

| (D) | Cluster 3 | |
|---|---|---|
| **Gene** | ***P* value** | **Average log2FC** |
| **MGST1** | 4.26E-19 | 2.243888 |
| **UBB** | 8.69E-18 | 1.853261 |
| **UCHL1** | 1.73E-38 | 3.566619 |
| **TRNP1** | 1.61E-20 | 1.883486 |
| **VIM** | 4.76E-30 | 4.172655 |
| **CKB** | 2.53E-19 | 1.844814 |
| **GLS** | 3.04E-28 | 1.925304 |
| **AREG** | 2.61E-16 | 1.895932 |
| **ODC1** | 1.16E-16 | 2.601342 |
| **SERPINE1** | 0.002905 | 2.408616 |

**Table 5:** Cell type annotation in hyperproliferative and hypoproliferative spheroids

**Related to Figure 7E and F**

| | Hyperproliferative spheroid (total scores of hit) | | | | Hypoproliferative spheroids (total scores of hit) | | | | | |
|---|---|---|---|---|---|---|---|---|---|---|
| **Cell type** | **Cluster 0** | **Cluster 1** | **Cluster 2** | **Cluster 3** | **Cluster 0** | **Cluster 1** | **Cluster 2** | **Cluster 3** | **Cluster 4** | **Cluster 5** |
| **Epithelial cell** | 1614.45 | 839.41 | 1405.52 | 235.7 | 750.5 | 851.35 | 2947.69 | 1204.55 | 0 | 0 |
| **Basal cell** | 387.95 | 2081.23 | 976.71 | 386.87 | 1162.92 | 1269.72 | 1050.41 | 0 | 0 | 443.24 |
| **Neoplastic cell** | 627.49 | **5515.99** | 480.22 | 950.37 | 1369.88 | 2722.49 | 692.01 | 550.15 | 0 | 313.83 |
| **Stem cell** | 0 | 951.92 | 0 | 0 | 221.66 | 632.08 | 0 | 0 | 0 | 0 |
| **Goblet cell** | 516.09 | 0 | 0 | 0 | 278.52 | 0 | 0 | 539.61 | 0 | 0 |
| **Club cell** | 338.83 | 0 | 0 | 0 | 0 | 0 | 0 | 0 | 0 | 0 |
| **Pneumocyte cell** | 176.72 | 0 | 0 | 0 | 0 | 0 | 0 | 0 | 0 | 0 |
| **Endothelial cell** | 134.11 | 0 | 0 | 0 | 0 | 0 | 0 | 0 | 0 | 0 |
| **Fibroblast cell** | 0 | 0 | 0 | 870.08 | 291.09 | 0 | 0 | 0 | 0 | 1791 |
| **Mesenchymal cell** | 0 | 0 | 0 | 114.06 | 0 | 457.31 | 0 | 0 | 0 | 0 |
| **Keratinizing barrrier epithelial cell** | 0 | 0 | 0 | 0 | 0 | 0 | 1121.49 | 0 | 0 | 0 |
| **Others cell** | 1134.33 | 0 | 0 | 0 | 0 | 0 | 0 | 0 | 0 | 0 |
