## Supplementary figures and images for "Clonal spheroids capture functional and genetic heterogeneity of head and neck cancer"

### Supplementary video

#### Slide 1
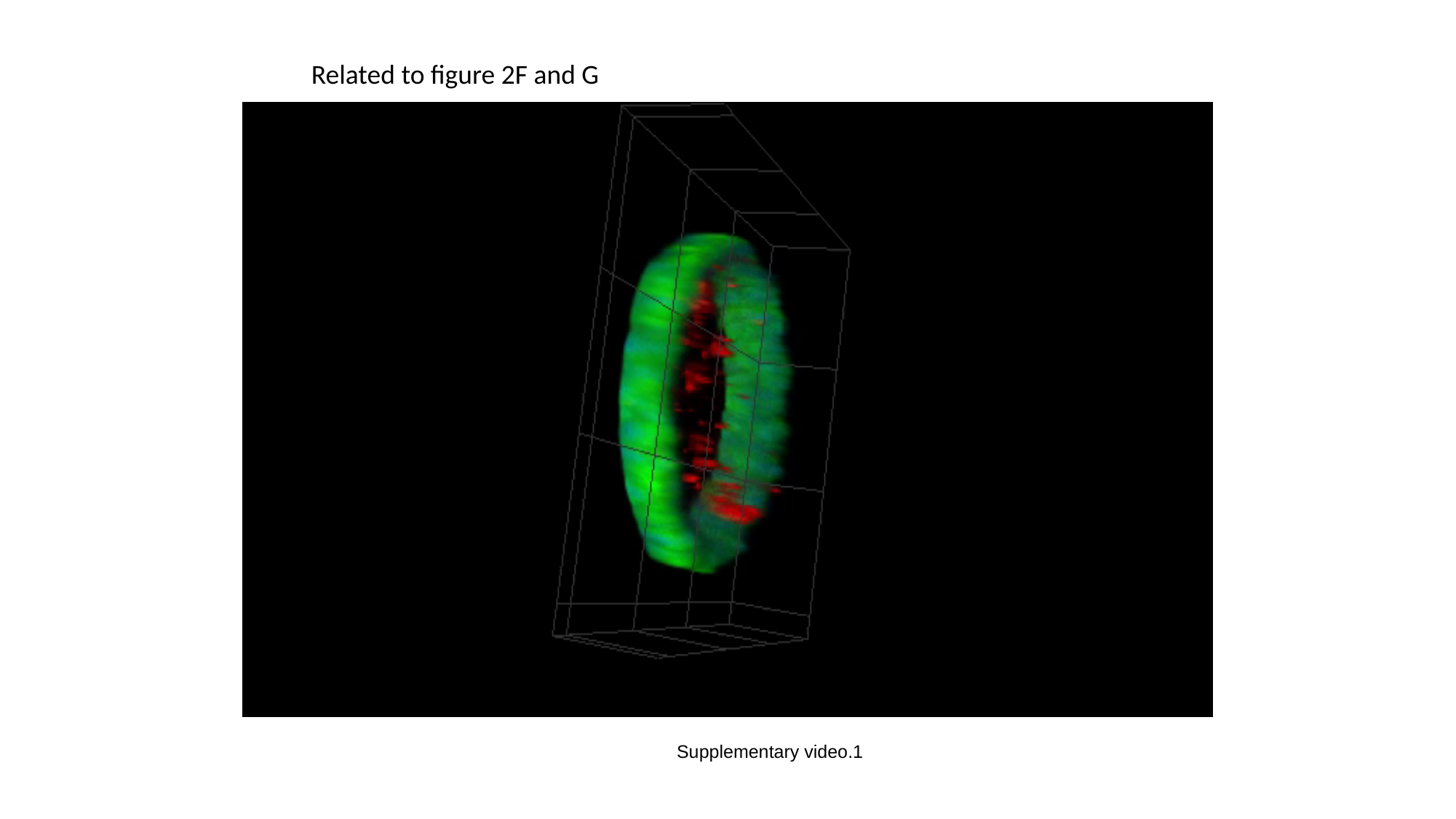

Related to figure 2F and G
Supplementary video.1
